# Data-driven models of dominantly-inherited Alzheimer’s disease progression

**DOI:** 10.1101/250654

**Authors:** Neil P. Oxtoby, Alexandra L. Young, David M. Cash, Tammie L. S. Benzinger, Anne M. Fagan, John C. Morris, Randall J. Bateman, Nick C. Fox, Jonathan M. Schott, Daniel C. Alexander

## Abstract

Dominantly-inherited Alzheimer’s disease is widely hoped to hold the key to developing interventions for sporadic late onset Alzheimer’s disease. We use emerging techniques in generative data-driven disease-progression modelling to characterise dominantly-inherited Alzheimer’s disease progression with unprecedented resolution, and without relying upon familial estimates of years until symptom onset (EYO). We retrospectively analysed biomarker data from the sixth data freeze of the Dominantly Inherited Alzheimer Network observational study, including measures of amyloid proteins and neurofibrillary tangles in the brain, regional brain volumes and cortical thicknesses, brain glucose hypometabolism, and cognitive performance from the Mini-Mental State Examination (all adjusted for age, years of education, sex, and head size, as appropriate). Data included 338 participants with known mutation status (211 mutation carriers: 163 *PSEN1*; 17 *PSEN2*; and 31 *APP*) and a baseline visit (age 19–66; up to four visits each, 1·1 ± 1·9 years in duration; spanning 30 years before, to 21 years after, parental age of symptom onset). We used an event-based model to estimate sequences of biomarker changes from baseline data across disease subtypes (mutation groups), and a differential-equation model to estimate biomarker trajectories from longitudinal data (up to 66 mutation carriers, all subtypes combined). The two models concur that biomarker abnormality proceeds as follows: amyloid deposition in cortical then sub-cortical regions (approximately 24±11 years before onset); CSF p-tau (17±8 years), tau and A*β*42 changes; neurodegeneration first in the putamen and nucleus accumbens (up to 6 ± 2 years); then cognitive decline (7 ± 6 years), cerebral hypometabolism (4 ± 4 years), and further regional neurodegeneration. Our models predicted symptom onset more accurately than EYO: root-mean-squared error of 1·35 years versus 5·54 years. The models reveal hidden detail on dominantly-inherited Alzheimer’s disease progression, as well as providing data-driven systems for fine-grained patient staging and prediction of symptom onset with great potential utility in clinical trials.

## 1. Introduction

Understanding and identifying the earliest pathological changes of Alzheimer’s disease is key to realizing disease-modifying treatments, which are likely to be most efficacious when given early. However, identifying individuals in the presymptomatic stage of typical, sporadic, late onset Alzheimer’s disease is challenging. Therefore, there is considerable interest in investigating dominantly-inherited Alzheimer’s disease, which is caused by mutations in the Amyloid Precursor Protein (*APP*), Presenilin 1 (*PSEN1*), and Presenilin 2 (*PSEN2*) genes, and which provides the opportunity to identify asymptomatic “at risk” individuals prior to the onset of cognitive decline for observational studies and clinical trials. Although considerably rarer than sporadic Alzheimer’s disease, dominantly-inherited Alzheimer’s disease has broadly similar clinical presentation (Tang et al., 2016, Ryan et al., 2016), i.e., episodic memory followed by further cognitive deficits, and both display heterogeneity in terms of symptoms and progression, much of which is unexplained (Bateman et al., 2010). An important question when attempting to extrapolate biomarker dynamics (and in due course clinical trials results) between dominantly-inherited Alzheimer’s disease and sporadic Alzheimer’s disease, is whether presymptomatic changes in dominantly-inherited Alzheimer’s disease mirror those in sporadic Alzheimer’s disease, as might be expected given the broad similarities in pathological features across both diseases (Bateman et al., 2010, Weiner et al., 2012, Morris et al., 2012, Cairns et al., 2015).

Most previous investigations into dominantly-inherited Alzheimer’s disease progression used traditional regression models to explore the time course of Alzheimer’s disease markers as a function of estimated years to onset (EYO) of clinical symptoms, based on age of onset (Ryman et al., 2014) in affected first-degree relatives. In 2012, this type of cross-sectional analysis of biomarker trajectories in the Dominantly Inherited Alzheimer Network (DIAN) observational study estimated the following sequence of pre-symptomatic biomarker changes (Bateman et al., 2012): measures of beta-amyloid (A*β*) in cerebrospinal fluid (CSF) and in amyloid imaging using Pittsburgh-compound B PET (PiB-PET); CSF levels of tau; regional brain atrophy; cortical glucose hypometabolism in flurodeoxyglucose PET (FDG-PET); episodic memory; Mini Mental State Examination (MMSE) score (Folstein et al., 1975); and Clinical Dementia Rating Sum of Boxes (CDRSUM) score (Berg, 1988). Results of a more recent model-based analysis (Fleisher et al., 2015) showed a similar progression sequence in the Alzheimer’s Prevention Initiative Colombian cohort, all of whom carry the same mutation (E280A PSEN1). In 2013, a more detailed investigation of imaging biomarkers (Benzinger et al., 2013) observed regional variability in the cross-sectional sequence of biomarker changes: some grey-matter structures having amyloid plaques may not later lose metabolic function, and others may not atrophy. Various other studies of dominantly-inherited Alzheimer’s disease have reported early behavioural changes (Ringman et al., 2015) and presymptomatic within-individual atrophy (Cash et al., 2013) in brain regions commonly associated with sporadic Alzheimer’s disease, and additionally in the putamen and thalamus. The key feature in each of these studies of dominantly-inherited Alzheimer’s disease progression is the reliance upon EYO, which is typically based upon the estimated age at which an individual’s affected parent first shows progressive cognitive decline (Bateman et al., 2010), or upon the average age of onset for a mutation type (Ryman et al., 2014). The parental estimate of familial age of onset is generated by a semi-structured interview and is known to be inherently uncertain both because of uncertainties in estimating when an individual is deemed to be affected, and because there can be substantial within-family and within-mutation differences in actual age of onset (Ryman et al., 2014). This uncertainty in EYO limits its utility for estimating disease progression in pre-symptomatic individuals who carry a dominantly-inherited Alzheimer’s disease mutation: reducing confidence in predicting onset; and when staging patients — at best reducing the resolution in which biomarker ordering can be inferred, at worst biasing the ordering. The mutation age of onset addresses higher variance estimates, and demonstrates good predictability, but is still a retrospective estimate and not obtained by prospectively monitoring for actual onset of symptoms in a uniform longitudinal fashion.

Here we take a different approach: generative, data-driven, disease progression modelling. Data-driven progression models have emerged in recent years as a family of computational approaches for analyzing progressive diseases. Instead of regressing against pre-defined disease stages (Bateman et al., 2012, Ridha et al., 2006, Scahill et al., 2002, Yang et al., 2011), or learning to classify cases from a labelled training database (Klöppel et al., 2008, Mattila et al., 2011, Young et al., 2013), generative data-driven progression models construct an explicit quantitative disease signature without the need for a priori staging. Mostly applied to neurodegenerative conditions like Alzheimer’s disease, results include discrete models of biomarker changes (Fonteijn et al., 2012, Young et al., 2014, Venkatraghavan et al., 2017), continuous models of biomarker dynamics (Jedynak et al., 2012, Villemagne et al., 2013, Donohue et al., 2014, Oxtoby et al., 2014), spatiotemporal models of brain image dynamics (Durrleman et al., 2013, Lorenzi et al., 2015, Schiratti et al., 2015, Huizinga et al., 2016), and models of disease propagation mechanisms (Seeley et al., 2009, Zhou et al., 2012, Raj et al., 2012, Iturria-Medina et al., 2014, 2017).

In this paper we use two generative data-driven disease progression models to extract patterns of observable biomarker changes in dominantly-inherited Alzheimer’s disease. We estimate ordered sequences of biomarker abnormality in disease subtypes (mutation groups) from cross-sectional data using an event-based model (Fonteijn et al., 2012, Young et al., 2014), and we estimate biomarker trajectories from short-interval longitudinal data using a nonparametric differential equation model similar to previous parametric work (Oxtoby et al., 2014, Villemagne et al., 2013). Our data-driven generative models have several potential advantages over previous models. First, they are generalizable to non-familial forms of progressive diseases because they do not rely on EYO. Second, they generate a uniquely detailed sequence of biomarker changes and trajectories. Third, they support a fine-grained staging and prognosis system of potential direct application to clinical trials and in clinical practice. We demonstrate the prognostic utility by predicting actual symptom onset in unseen data more accurately than using EYO.

## 2. Materials and Methods

We used data-driven models to analyse biomarker data (MRI, PET, CSF, cognitive test scores) from the DIAN study. From cross-sectional (baseline) data we estimated disease progression sequences using an event-based model (Fonteijn et al., 2012, Young et al., 2014). For explicit quantification of disease progression time, we estimated biomarker trajectories from short-term longitudinal data by using covariate-adjusted, nonparametric differential equation models, which offer two key advantages over previous approaches in (Villemagne et al., 2013, Oxtoby et al., 2014): replacing parametric model-selection with a data-driven approach, and explicitly estimating population variance in a Bayesian manner. Models were cross-validated internally (see Statistical Analysis section and Supplementary Material).

### 2.1. Participants

At the sixth data freeze, the DIAN cohort included 338 individual participants (192 females, 57%) with known mutation status and a baseline visit, aged 19–66 years at baseline (39±10 years), with up to four visits each (1·1 ± 1·9 years in duration, total of 535 visits), spanning 30 years before and 21 years after parental age of symptom onset. For detailed descriptive summaries of the DIAN cohort, we refer the reader to Morris et al. (2012).

#### 2.1.1. Data selection and preparation

Table 1 summarises the demographics of DIAN participants analysed in this work.

We selected 24 Alzheimer’s disease biomarkers based on specificity to the disease, or if disease ‘signal’ is present, i.e., quantifiable distinction between mutation carriers and non-carriers (see Statistical Analysis section). The biomarkers include CSF measures of molecular pathology (amyloid proteins and neurofibrillary tangles); a cognitive test score (MMSE); regional brain volumetry from MRI, e.g., hippocampus, middle-temporal region, temporo-parietal cortex; PiB-PET imaging measures of amyloid accumulation; and FDG-PET imaging measures of glucose hypometabolism. We excluded imaging data (21 structural scans from 10 participants) having artefacts or non-Alzheimer’s disease pathology such as a brain tumour. Of the included participants, 211 (117 females, 55%) were dominantly-inherited Alzheimer’s disease mutation carriers (MC): 163 *PSEN1*; 17 *PSEN2*; and 31 *APP*; and 120 were non-carriers (NC). Baseline data for mutation carriers and non-carriers was used to fit event-based models. The full set of biomarkers included in the event-based model is listed on the vertical axis of Figure 1.

Of the 211 included mutation carriers, 66 had longitudinal data suitable for fitting differential equation models. To reduce the influence of undue measurement noise we excluded biomarker data with a large coefficient of variation within individuals, e.g., as done for CSF biomarkers in Bateman et al. (2012). Following Villemagne et al. (2013), we also excluded differential data that was both normal (beyond a threshold determined by clustering), and non-progressing (rate of change has a contradictory sign to disease progression, e.g., reverse atrophy or improved cognition). Finally, we identified six cognitively normal mutation carriers who developed symptoms during the study (global CDR becoming nonzero after baseline). Since we use our differential equation models to predict symptom onset for these participants, we excluded them from the model fits to avoid circularity (including them doesn’t alter our results considerably). This left data from up to 51 mutation carriers (41 *PSEN1*; 1 *PSEN2*; 9 *APP*; 28 females) available for analysis using differential equation models. Subsets had data for structural imaging (47; 26 females), CSF (32; 16 females), PiB PET (32; 16 females), and FDG PET (39; 22 females) biomarkers.

We used stepwise regression to remove the influence of age, years of education, sex, and head size (total intracranial volume, MRI volumes only) prior to fitting our models.

### 2.2. Models

#### 2.2.1. Cross-sectional: Event-Based Models

The event-based model infers a sequence in which biomarkers show abnormality, together with uncertainty in that sequence, from cross-sectional data (Fonteijn et al., 2012). This longitudinal picture of disease progression is estimable using this approach because, across the spectrum of Dominantly Inherited Alzheimer Network study participants from cognitively normal controls (non-carriers of dominantly-inherited Alzheimer’s disease mutations), to presymptomatic mutation carriers, and symptomatic patients, more individuals will show higher likelihood of abnormality in biomarkers that change early in the progression. Thus, with sufficient representation across combinations of abnormal and normal observations, the likelihood of any full ordered sequence can be estimated to reveal the most likely sequences. The probabilistic sequence of events estimated by the event-based model is useful for fine-grained staging of individuals by calculating the likelihood of their data (biomarker observations) arising from each stage of the sequence (Young et al., 2014).

**Table 1:** Demographics for DIAN participants at Data Freeze 6. **A.** Cross-sectional data used to build event-based models of dominantly-inherited Alzheimer’s disease progression. **B.** Longitudinal data used to build differential equation models of dominantly-inherited Alzheimer’s disease progression. See Data Selection and Preparation section for more. Abbreviations: Cog – Cognitive test scores; PiB – Pittsburgh compound B PET data; FDG – fludeoxyglucose hypometabolism PET data; EYO – estimated years to onset based on parental age of symptom onset.

| Demographic |  | Non-carriers | Mutation carriers [*PSEN1*, *PSEN2*, *APP*] |
| --- | --- | --- | --- |
| **A.** | *n* analysed, cross-sectional (event-based models) | 127 | 211 [163, 17, 31 (77%, 8%, 15%)] |
|  | Female | 75 (59%) | 117 [92, 5, 20 (79%, 4%, 17%)] |
|  | APOE4-positive | 37 (29%) | 61 [47, 7, 7 (77%, 11.5%, 11.5%)] |
|  | APOE4-negative | 90 (71%) | 150 [116, 10, 24 (77%, 7%, 16%)] |
| | Age at baseline ± standard deviation, years | 39 ± 10 | 39 ± 10 [39 ± 10, 39 ± 10, 43 ± 10] |
| | Education at baseline ± standard deviation, years | 15 ± 3 | 14 ± 3 [14 ± 3, 15 ± 3, 14 ± 3] |
| | EYO at baseline ± standard deviation, years | -7 ± 12 | -7 ± 10 [-7 ± 10, -12 ± 10, -6 ± 9] |
| **B.** | *n* analysed, longitudinal (differential equation models) | n/a | Cog: 51 [41, 1, 9 (80%, 2%, 18%)]<br>MRI: 47 [37, 2, 8 (79%, 4%, 17%)]<br>CSF: 32 [28, 1, 3 (88%, 3%, 9%)]<br>PiB: 32 [25, 2, 5 (78%, 6%, 16%)]<br>FDG: 39 [31, 2, 6 (80%, 5%, 15%)] |
|  | Female | n/a | Cog: 28 (55%) [21, 1, 6 (75%, 4%, 21%)]<br>MRI: 26 (55%) [19, 1, 6 (73%, 4%, 23%)]<br>CSF: 16 (50%) [13, 0, 3 (81%, 0%, 19%)]<br>PiB: 16 (50%) [11, 1, 4 (69%, 6%, 25%)]<br>FDG: 22 (56%) [16, 1, 5 (73%, 5%, 23%)] |
|  | APOE4-positive | n/a | Cog: 17 (33%) [13, 0, 4 (76%, 0%, 24%)]<br>MRI: 16 (34%) [11, 1, 4 (69%, 6%, 25%)]<br>CSF: 8 (25%) [6, 1, 1 (75%, 12.5%, 12.5%)]<br>PiB: 13 (41%) [9, 1, 3 (69%, 8%, 23%)]<br>FDG: 14 (36%) [10, 1, 3 (71%, 7%, 21%)] |
|  | APOE4-negative | n/a | Cog: 34 (67%) [28, 1, 5 (82%, 3%, 15%)]<br>MRI: 31 (66%) [26, 1, 4 (84%, 3%, 13%)]<br>CSF: 24 (75%) [22, 0, 2 (92%, 0%, 8%)]<br>PiB: 19 (59%) [16, 1, 2 (84%, 5%, 11%)]<br>FDG: 25 (64%) [21, 1, 3 (84%, 4%, 12%)] |
| | Age at baseline ± standard deviation, years | n/a | Cog: 41 ± 10 [40 ± 10, 32 ± 0, 48 ± 7]<br>MRI: 42 ± 10 [40 ± 10, 45 ± 18, 50 ± 6]<br>CSF: 43 ± 9 [42 ± 9, 57 ± 0, 48 ± 8]<br>PiB: 43 ± 10 [41 ± 10, 45 ± 18, 49 ± 4]<br>FDG: 42 ± 10 [41 ± 10, 45 ± 18, 48 ± 5] |
| | Education at baseline ± standard deviation, years | n/a | Cog: 14 ± 2 [14 ± 2, 18 ± 0, 15 ± 2]<br>MRI: 14 ± 2 [14 ± 2, 15 ± 4, 15 ± 2]<br>CSF: 14 ± 3 [14 ± 3, 12 ± 0, 14 ± 3]<br>PiB: 14 ± 3 [14 ± 2, 15 ± 4, 15 ± 3]<br>FDG: 14 ± 2 [14 ± 2, 15 ± 4, 15 ± 2] |
| | EYO at baseline ± standard deviation, years | n/a | Cog: -3 ± 7 [-3 ± 7, -19 ± 0, -2 ± 8]<br>MRI: -2 ± 8 [-3 ± 7, -6 ± 18, 0 ± 6]<br>CSF: 0 ± 7 [0 ± 7, 7 ± 0, -3 ± 6]<br>PiB: -2 ± 7 [-3 ± 6, -6 ± 18, 0 ± 3]<br>FDG: -3 ± 7 [-4 ± 7, -6 ± 18, -2 ± 5] |

We fit an event-based model to determine the most probable sequence of biomarker abnormality events and the uncertainty in this sequence for all but 3 of the 24 biomarkers described previously: measurements of entorhinal cortex, thalamus, and caudate volume were excluded on the basis that they did not show significant differences (see Statistical Analysis section) between non-carriers and symptomatic mutation carriers after correction for age, sex, education and total intracranial volume. Each event represents the transition of a biomarker from a normal level (as seen in non-carriers) to an abnormal level (as seen in symptomatic patients). The probability a biomarker measurement is normal is modelled as a Gaussian distribution, and estimated using data from non-carriers. The distribution of abnormal measurements is also modelled as a Gaussian distribution, but estimated by fitting a mixture of two Gaussians (Fonteijn et al., 2012) to data from all mutation carriers: the first Gaussian models the distribution of normal measurements, and is kept fixed to the values estimated from non-carriers; the second Gaussian models the distribution of abnormal measurements, and is optimised using data from mutation carriers. The sequence of events was estimated in various population subgroups: all 211 mutation carriers; 163 *PSEN1* mutation carriers; 17 *PSEN2* mutation carriers; and 31 *APP* mutation carriers. We also considered separate event-based models by APOE4 status: 61 mutation carriers who were APOE4-positive (with one or more APOE4 alleles), and 150 mutation carriers who were APOE4-negative. For further details of the model fitting procedures, see the Statistical Analysis section. We assigned participants to patient stages based on their most probable position along the most probable event sequence (Young et al., 2014) for all mutation carriers combined. We assessed the efficacy of the patient staging system using only participants with data available for all biomarkers (*n* = 30, total of 42 follow-up visits), as missing entries cause uncertainty in a participant’s model stage.

#### 2.2.2. Longitudinal: Differential Equation Models

Reconstruction of biomarker trajectories ideally requires dense longitudinal data collected over the full time course of the disease. Such data is not yet available due to the prohibitive expense and complexity of collection, which means that we must resort to alternative methods. In dominantly-inherited Alzheimer’s disease and other neurodegenerative diseases, the availability of short-term longitudinal data of a few years permits estimation of an individual’s rate of change over that time span, e.g., via linear regression. These short-interval longitudinal observations are interpreted as noisy samples (segments) from an average biomarker trajectory. Instead of attempting to align the raw data segments, e.g., (Donohue et al., 2014), they are used to generate a cross-section of *di*ff*erential* data: biomarker rate-of-change as a function of biomarker value, i.e., a differential equation. For sufficient coverage across a range of biomarker values tracking dominantly-inherited Alzheimer’s disease progression, the function can be integrated to produce a trajectory. The technique estimates probabilistic trajectories that are useful for predicting symptom onset (with confidence bounds) for an individual by aligning their biomarker observations to the trajectory. We refer to this approach as a differential equation model, since it is analogous to the numerical integration of ordinary differential equations as performed in many fields of science.

Here we took a univariate approach, considering each biomarker in turn. We found each individual’s biomarker rate-of-change, *dx*/*dt*, through linear regression of their short-term longitudinal data. We then plotted this against average biomarker value *x* (for each individual), and fit a curve *f* (*x*)

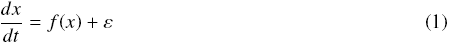

with residual errors *ε*. Finally, we numerically integrated along the fitted DEM curve *C* between biomarker extreme values *a* and *b* to estimate a model of the average trajectory

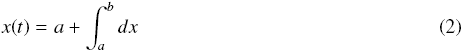

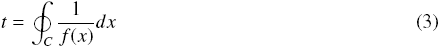

Previously, Villemagne et al. (2013) took a model-selection approach to finding the best fitting polynomial regression model for *f* (*x*). Instead, we use a nonparametric Bayesian approach (see Statistical Analysis section).

In dominantly-inherited Alzheimer’s disease, biomarker trajectories are usually investigated as a function of EYO. This is an approximate proxy for disease progression time where EYO = 0 is the estimated point of onset of clinical symptoms, based on familial age of onset such as that of an affected parent. Here we defined *t* = 0 at a data-driven canonical abnormal level: the median biomarker value for symptomatic participants in the DIAN cohort (first symptomatic visit).

A quantity of clinical interest is the interval of time between normal and abnormal biomarker levels, which we refer to as the Abnormality Transition Time, and define in a data-driven manner via median values for asymptomatic (canonically normal) and symptomatic (canonically abnormal as above) participants in the DIAN dataset. Our probabilistic approach produces an abnormality transition time *distribution* per biomarker. The cumulative probability of abnormality produces data-driven sigmoid-like curves, which we combine across biomarkers to estimate a temporal pattern of disease progression.

We estimated time to onset (ETO) for each individual as a weighted average across biomarkers. Aligning each biomarker measurement to each biomarker trajectory produces a set of biomarker-specific ETOs with credible intervals. We weighted by the inverse width of the credible interval, to assign lower confidence to estimates with large uncertainty. Incomplete data was used, with missing values omitted from the weighted average.

### 2.3. Statistical Analysis

For fitting the event-based model we followed the same procedures as in Young et al. (2014). Briefly, the characteristic sequence and its uncertainty is estimated through a Markov chain Monte Carlo sampling procedure with greedy-ascent initialisation for maximising the data likelihood (Fonteijn et al., 2012). We used a noninformative uniform prior on the sequence. When fitting an event-based model, it is important to select a set of biomarkers specific to the disease. That is, where disease ‘signal’ is present: a quantifiable distinction between normal and abnormal. For this procedure we used a paired t-test, and thresholded significance at *p* < 0.01/24, Bonferroni-corrected for multiple comparisons. We accounted for missing data by imputing biomarker values such that missing measurements had an equal probability of being normal or abnormal (Young et al., 2015), and thus would not influence the population sequence. We performed cross-validation of the event-based model by re-estimating the event distributions and maximum likelihood sequence for 100 bootstrap samples from each data subset. The positional variance diagrams for the cross validation results show the proportion of bootstrap samples in which event *i* (vertical axis) appears at position *k* (horizontal axis) of the maximum likelihood sequence.

For fitting differential equation models we use a nonparametric approach known as Gaussian process regression (Rasmussen and Williams, 2006) to produce a probabilistic fit (a distribution of curves) that is determined by the data. The fitting was implemented within the probabilistic programming language Stan (Stan Development Team, 2015), which performs full Bayesian statistical inference using Markov chain Monte Carlo sampling and penalized maximum likelihood estimation. We used a vanilla squared-exponential kernel (Rasmussen and Williams, 2006) for the Gaussian process prior covariance: 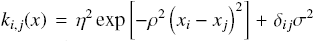 with hyperparameters η, ρ, σ and Kronecker delta function δ. The Gaussian process prior hyperparameters guide the shape of the regression function, *f* (*x*), and were also estimated from the data. Here we used weakly-informative broad half-Cauchy hyperparameter priors, and diffuse initial conditions to aid model identifiability. To check for over-fitting we performed 10-fold cross-validation (see Supplementary Material), and various posterior predictive checks to assess model quality and numerical convergence (Gelman et al., 2014, Vehtari et al., 2016).

## 3. Results

We first present our cross-sectional multimodal modelling of the fine-grained ordering of dominantly-inherited Alzheimer’s disease biomarker abnormality using an event-based model. We then present our longitudinal modelling of dominantly-inherited Alzheimer’s disease biomarker trajectories using differential equation models.

### 3.1. Cross-sectional results: event-based models of biomarker abnormality sequences

Figure 1 is a positional variance diagram of the maximum likelihood sequence of biomarker abnormality events (top to bottom), and its uncertainty (left to right), across all available 211 mutation carriers in the DIAN dataset. Grayscale intensity represents confidence in each event’s position within the sequence, and is calculated from Markov chain Monte Carlo samples from the event-based model (Young et al., 2014). The closer this diagram is to a black diagonal, the more confidence there is in the disease progression sequence.

The event-based model reveals a distinct sequence of biomarker abnormality in dominantly-inherited Alzheimer’s disease: regional (cortical then striatal) amyloid deposition on PiB-PET scans; CSF measures of neuronal injury (total tau), neurofibrillary tangles (phosphorylated tau), and amyloid plaques (A*β*42 and A*β*40/A*β*42 ratio); MRI measures of volume loss in the putamen and nucleus accumbens; global cognition (MMSE score). Thereafter the ordering in which FDGPET hypometabolism and other MRI measures become abnormal is less certain. We found high certainty early in the ordering of these biomarkers (as reflected by the more solid blocks along the diagonal), with lower certainty later in the ordering of regional volumes (more diffuse grey blocks straying from the diagonal). This pattern (left) persists under cross-validation (right).

We also fit the event-based model to subgroups of the mutation carriers in the DIAN data set. Figure 2 shows positional variance diagrams of the biomarker abnormality event sequence in APOE4-positive and APOE4-negative participants (those with and without an apolipoprotein-4 allele), and for the three dominantly-inherited Alzheimer’s disease mutation types in DIAN: *PSEN1*, *PSEN2*, and *APP*. For ease of comparison, the sequence ordering on the vertical axes of each plot was chosen to be the most probable ordering from Figure 1 (all mutation carriers). Cross validation results for Figure 2 are shown in Supplementary Figure 1.

Broadly speaking, we see good agreement of the event sequences across APOE4 subgroups and across mutation type subgroups in Figure 2, with some subtle differences between groups: earlier CSF A*β*42 and A*β*40/A*β*42 ratio in the APOE4-positive and *APP* groups; earlier volume abnormality in the fusiform gyrus for the *PSEN2* group; earlier putamen volume abnormality for the *APP* group. The uncertainty is high in the subgroup orderings due in part to the low numbers of participants in these groups, which reduces power to draw concrete conclusions based on these subtle differences between groups.

**Figure 1:**
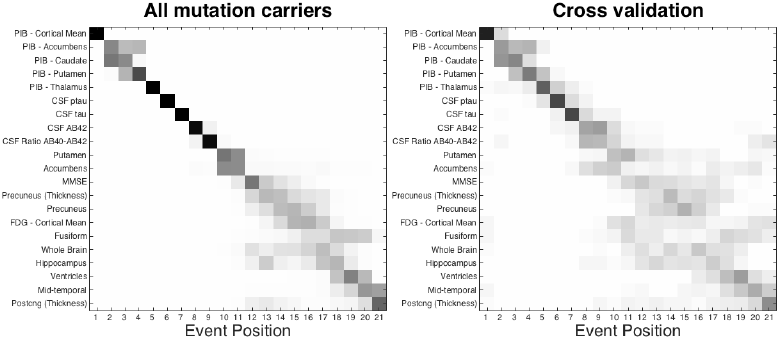
Event-based model progression of dominantly-inherited Alzheimer’s disease progression. Positional variance diagrams. Left: event-based model estimated on all mutation carriers in the DIAN dataset. Right: cross-validation through bootstrapping. The vertical ordering (top to bottom) is given by the maximum likelihood sequence estimated by the model. Grayscale intensity represents posterior confidence in each event’s position (each row), from Markov chain Monte Carlo samples of the posterior.

Figure 3 demonstrates the fine-grained staging capabilities of the event-based model. Using the model for all mutation types (Figure 1), each participant in the DIAN dataset was assigned a disease stage that best reflects their measurements (see Materials and Methods section, and Young et al. (2014)). The staging proportions are shown in Figure 3(a), differentiated by broad diagnostic groups (CN: Cognitively Normal, global CDR = 0; MCI: very mild dementia consistent with Mild Cognitive Impairment, global CDR = 0·5; AD: probable dementia due to Alzheimer’s disease, global CDR > 0·5). Longitudinal consistency of staging is shown in Figure 3(b) where each participant’s baseline stage is plotted against available follow-up stages between baseline and months 12/24/36.

The baseline staging in Figure 3(a) shows good separation of diagnostic groups: all of the non-carriers are assigned to stage 0 (black), CN mutation carriers (green) are at earlier model stages, mutation carriers diagnosed with probable Alzheimer’s disease dementia (red) are at late model stages, and mutation carriers with very mild dementia consistent with MCI (blue) are more spread out across the stages. Within carriers, the model shows high classification accuracy for separating those who are cognitively normal from those with probable dementia: a balanced accuracy of 93% is achieved by classifying participants above stage 10 (putamen volume abnormality) as having probable Alzheimer’s disease dementia. This shows that our generative model can also be used for discriminative applications with performance comparable to state-of-the-art multimodal binary classifiers (Willette et al., 2014). Further, Supplementary Figure 15(a) shows positive associations between baseline EYO and baseline event-based model stage, by diagnostic group.

The follow-up staging in Figure 3(b) shows good longitudinal consistency: at 32 of 36 (89%) follow-up time points the model stage is the same or it increased; at 34 of 36 (94%) follow-up time points the stage was either unchanged, it increased, or it decreased within the uncertainty of the ordering. This included the clinical converters, which are shown with triangles (CN to MCI in green; MCI to AD in red). The remaining two follow-up time points at which the model stage decreased (green circles in Figure 3(b); 1 *PSEN1*, 1 *APP*) have inconsistent amyloid levels between CSF and regional PiB-PET, potentially due to discord between these biomarkers as has been observed in some individuals (Landau et al., 2013, Schroeter et al., 2015).

**Figure 2:**
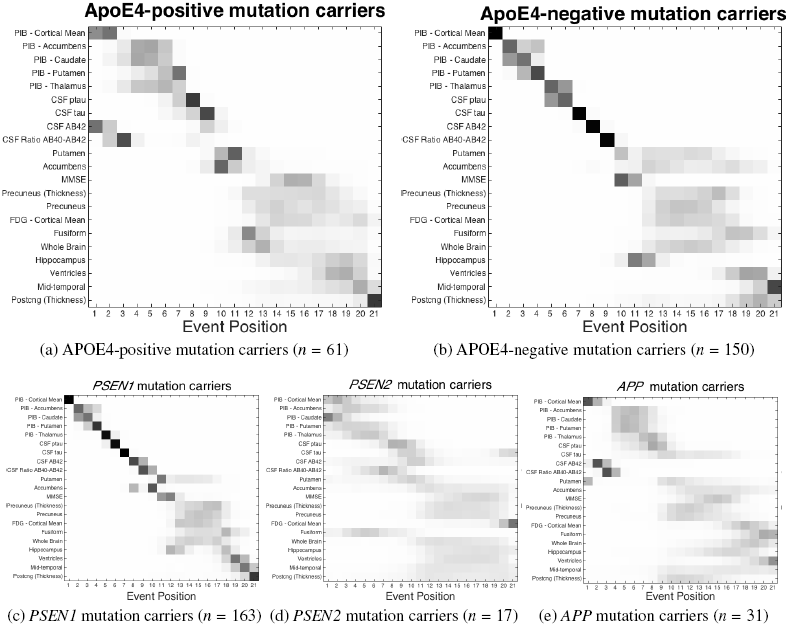
Event-based models of dominantly-inherited Alzheimer’s disease: mutation groups. Data-driven sequences of biomarker abnormality shown as positional variance diagrams for mutation carriers in the DIAN dataset who are (a) APOE4-positive; (b) APOE4-negative; (c) *PSEN1* mutation carriers; (d) *PSEN2* mutation carriers; (e) *APP* mutation carriers. Compare with Figure 1 (all groups combined): similar ordering, with a notable difference: APOE4-positive participants and *APP* mutation carriers both showed earlier CSF A*β*42 abnormality. See Supplementary Figure 1 for cross-validation results.

### 3.2. Longitudinal results: biomarker trajectories from differential equation models

Figure 4 shows a selection of dominantly-inherited Alzheimer’s disease biomarker trajectories estimated from the DIAN dataset using our approach. Each average trajectory is shown as a heavy dashed black line, with uncertainty indicated by thin grey trajectories sampled from the posterior distribution. The time axis is defined such that *t* = 0 corresponds to the median biomarker value for symptomatic mutation carriers (sMC) in the DIAN dataset, which we define as the canonical abnormal level. This is marked in Figure 4 by a red horizontal line for each biomarker, with the corresponding distribution of biomarker values for sMC shown to the left of each trajectory as a red quartile box plot. The green quartile box plots show the biomarker distributions for asymptomatic mutation carriers (aMC), with the median value for each biomarker defining our canonical normal level and shown by a green horizontal line. Importantly, the canonical normal and abnormal levels are not required to estimate the biomarker trajectories, but are used to define the Abnormality Transition Time for each biomarker as a data-driven estimate of the duration of the transition between these levels. Our Bayesian approach estimates an Abnormality Transition Time density (probability distribution) for each biomarker, which is shown in blue in Figure 4. For comparison, linear mixed model fits to baseline DIAN data from Bateman et al. (2012) are shown in magenta in Figure 4 (b)/(c)/(d)/(e) (others not available).

**Figure 3:**
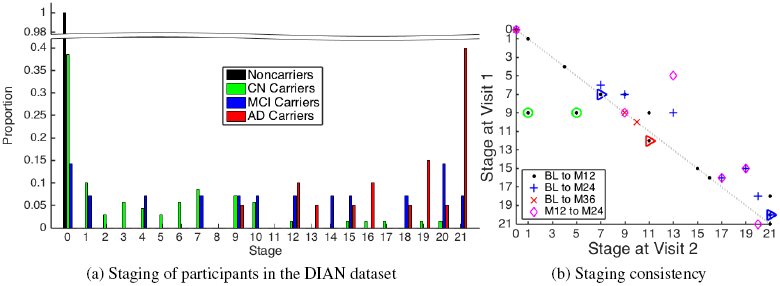
Event-based model staging results for dominantly-inherited Alzheimer’s disease. (a) Proportions grouped by diagnostic group: all noncarriers are at stage zero (black), and advancing disease stage is correlated strongly with cognitive impairment (green to blue to red). (b) Staging consistency across visits within three years of baseline for the *n* = 30 participants having complete longitudinal data (18 mutation carriers; 16 *PSEN1*, 2 *APP*). Most participants advance to a later stage (disease progresses towards the right). Green circles show the two participants (1 *PSEN1*, 1 *APP*) who regressed to earlier event-based model stages, which arises in both cases due to discordant amyloid measurements between CSF and PiB-PET. The triangles indicate clinical progressors (CN to MCI in blue; MCI to AD in red). BL: baseline; M: month; CN: Cognitively Normal (global CDR = 0); MCI: very mild dementia consistent with Mild Cognitive Impairment (global CDR = 0·5); AD: probable dementia due to dominantly-inherited Alzheimer’s disease (global CDR > 0·5).

Most trajectories in Figure 4 (and in Supplementary Material) show acceleration from normal to abnormal levels, with little evidence for post-onset deceleration/plateauing that would be consistent with the sigmoidal behaviour hypothesized for sporadic Alzheimer’s disease in, e.g., Jack et al. (2010). Biomarkers with trajectories that do not plateau, but remain dynamic into the symptomatic phase of the disease, offer potential utility for monitoring progression later in the disease. The grey curves capture uncertainty in the biomarker dynamics which arises both from fitting the differential equation models to discrete data, and from heterogeneity in the population. For comparison with our data-driven approach, the magenta trajectories in Figure 4 (b)/(c)/(d)/(e) are from Bateman et al. (2012), which used regression of baseline data against EYO. Qualitatively, they broadly agree with our trajectories for PiB-PET (cortical average amyloid deposition), MMSE, Hippocampus volume, and FDG-PET (cortical average hypometabolism), although, around symptom onset and beyond, our steeper Hippocampus volume trajectory implies a more aggressive progression than estimated cross-sectionally in Bateman et al. (2012).

**Figure 4:**
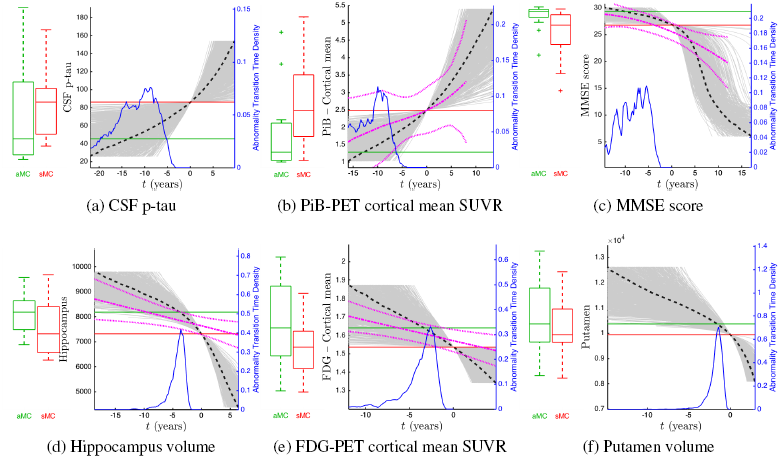
Differential equation models: dominantly-inherited Alzheimer’s disease biomarker trajectories. Shown are fits for selected biomarkers (see Models section). Fits for other biomarkers are provided in the Supplementary Material. Heavy black dashed lines show the average trajectory, with grey lines showing trajectories sampled from the posterior distribution. Time is expressed relative to the median biomarker value (red line) for symptomatic mutation carriers in the DIAN dataset (first visit with a nonzero CDR score), so that negative time suggests the average pre-symptomatic phase of dominantly-inherited Alzheimer’s disease. Box plots show biomarker distributions for asymptomatic (green, left, canonical normal) and symptomatic (red, right, canonical abnormal) mutation carriers (denoted aMC and sMC, respectively), with the distribution for estimated time between canonical normal and canonical abnormal (Abnormality Transition Time) shown in blue. Details of included participants are given in Table 1. For comparison, the magenta fits in (b), (c), (d), and (e) are those from the linear mixed models of baseline DIAN data against EYO from Bateman et al. (2012). SUVR = standardized uptake value ratio (relative to the cerebellum).

Figure 5 shows the cumulative probability for each biomarker in Figure 4. That is, the empirical distribution function for the Abnormality Transition Time densities in Figure 4, using the same time axis but on a logarithmic scale to ease visualisation. From Figure 5 we can infer an ordering of abnormality (Figure 5 legend) by comparing the times at which each curve reaches an abnormality probability of 0·5.

The cumulative probability curves in Figure 5 give a sense of both the average temporal ordering of biomarker abnormality (relative location of the curves at probability = 0·5), and the rate of progression (curve steepness) in the pre-symptomatic phase of dominantly-inherited Alzheimer’s disease. Whereas the event-based model approach is explicitly designed to infer an ordered sequence, our differential equation model approach is not. Nonetheless, the curves bear some resemblance to the hypothetical model in Jack et al. (2010), with the earliest phase of preclinical disease showing dynamic molecular pathology (CSF p-tau and PiB-PET), and other biomarkers becoming dynamic as onset approaches: cognitive decline (MMSE), neurodegeneration (MRI volumes), and hypometabolism (FDG-PET).

**Figure 5:**
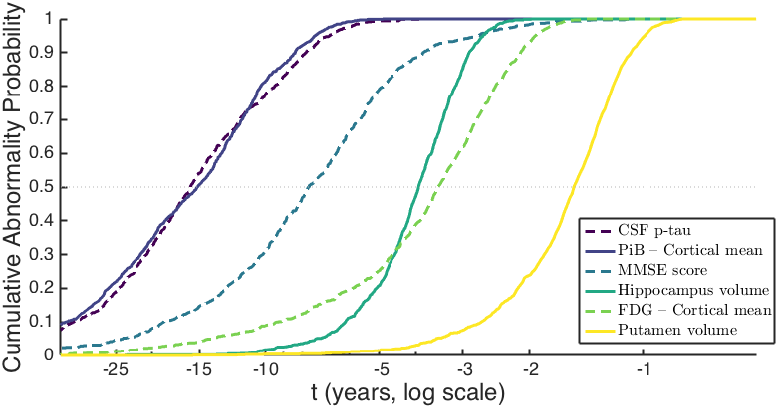
Differential equation models: selected data-driven sigmoids for dominantly-inherited Alzheimer’s disease biomarker progression. Cumulative probability of abnormality (vertical axis) is the empirical distribution of the Abnormality Transition Time in years prior to canonical abnormality (horizontal axis) as per Figure 4, calculated from each biomarker trajectory in Figure 4. The horizontal axis shows years prior to canonical abnormality. The order of biomarkers in the legend follows the order in which they reach a cumulative probability of abnormality of 0·5 (horizontal dotted grey line). Green-Blue-Yellow colour scale (viridis) with alternating solid/dashed lines in order of cumulative abnormality probability reaching 0·5 (legend).

### 3.3. Predicting time to symptom onset for unseen data

The models such as in Figure 4 further support an Estimated Time from Onset ETO*_m_* (together with uncertainty) for each biomarker *m* by aligning baseline biomarker measurement to the average trajectory. Uncertainty in ETO*_m_* is estimated from the corresponding probabilistic trajectory distribution (grey curves in Figure 4). A single estimate for each participant’s personal ETO, combining information from all biomarkers, then comes from averaging the ETO*_m_*, weighted by inverse uncertainty (here we used median absolute deviation in ETO*_m_*). For validation, we compare ETO to known actual years from onset for the six mutation carriers in the DIAN dataset who developed symptoms during the study (global CDR score becoming nonzero after baseline). These participants were omitted from the original differential equation model fits to avoid circularity.

Figure 6(a) plots estimated years from onset against actual years from onset for our model-derived ETO (red asterisks; dashed line fit), and for EYO (blue triangles; solid line fit), based on familial age of onset reported for an affected parent. A light grey line of reference shows perfect correspondence. Figure 6(b) shows quartile boxplots of the actual errors in predicting years to onset using ETO and EYO, at the visit where progression occurred.

The linear fits in Figure 6 (a) and boxplots in Figure 6(b) show that our data-driven ETO is a good predictor of actual years to onset with a root-mean-squared error of 1·34 years and a coefficient of determination of *r*^2^ ≈ 0·49. Familial age of onset is not as good: root-mean-squared error of 5·54 years and *r*^2^ ≈ 0·37 (EYO, based on parental age of onset); root-mean-squared error of 8·61 years and *r*^2^ ≈ 0·33 (Mutation EYO, based on mutation type). This poor performance is primarily because of very poor prediction for the participant at 3 years from onset (green circles, *PSEN1*). It is apparent from Figure 6 that our ETO may tend to overestimate when onset will occur (predicting earlier onset), and EYO tends to underestimate it (predicting later onset). This warrants further investigation with more data, but since onset may occur between visits to the clinic (interval censoring), it is likely more accurate to predict earlier onset, as our approach does.

**Figure 6:**
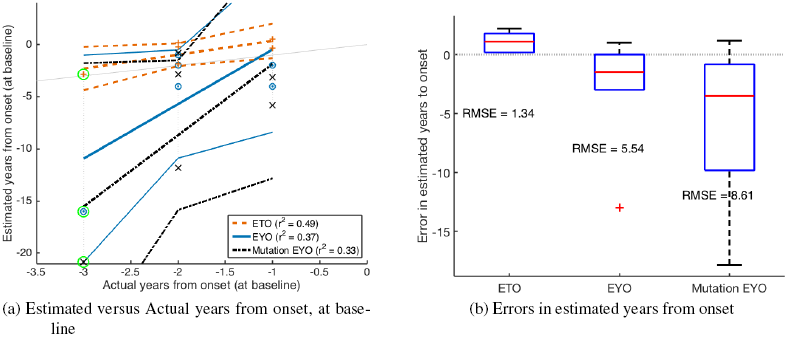
Predicting onset of clinical symptoms. For the six DIAN participants for whom global CDR became nonzero during the study (as of Data Freeze 6): (a) Estimated versus Actual years to onset at baseline using our model-based approach and using familial age of onset (EYO) and mutation type age of onset (Mutation EYO). Mutation EYO is calculated from the average age of onset within the three mutation types, using data from Table e-1 in Ryman et al. (2014), with the average weighted by the number of affected individuals per mutation. The light grey line shows perfect correlation as a reference and participants data points are connected by dotted grey vertical lines. Our model-derived ETO (red asterisks and dashed line fit) correlates with actual years to onset better than familial EYO (blue triangles and solid line fits), as shown by the adjusted coefficient of determination (*r*^2^). The green circle highlights an individual for whom our approach (ETO) is superior to the traditional approach (EYO) for predicting years to onset. (b) Quartile boxplots of the error in predicting onset using each estimate: ETO (left) has a superior root-mean-squared error (RMSE) to both EYO (middle) and Mutation EYO (right), and predicts symptom onset to occur sooner rather than later, which is likely to be more accurate due to interval censoring (symptom onset occurring between visits to the clinic).

### 3.4. Overview of results

Figure 7 visualises consistency across our two data-driven biomarker modelling approaches by showing patterns of dominantly-inherited Alzheimer’s disease progression obtained from each method on the DIAN dataset. The event-based model infers a probabilistic ordering of biomarker abnormality events through comparison of a cross-section of multi-modal observations, as shown for all mutation carriers in Figure 7(a) (reproduced from Figure 1). In contrast, each differential equation model works on an individual biomarker to estimate the biomarker trajectory. Figure 7(b) shows an alternative visualization of data-driven sigmoids for all included biomarkers, with the ordering determined as in Figure 5 by cumulative abnormality probability reaching 0·5 (black asterisks; white bars indicate the speed of biomarker change see caption of Figure 7 for details).

Qualitatively, Figure 7 shows that the different approaches estimate similar patterns of dominantly-inherited Alzheimer’s disease progression: accumulation of molecular pathology (amyloid, and tau where measured) followed by a blurring of cognitive abnormalities, brain hypometabolism, and regional changes to brain volume and cortical thickness. The combination of both models enables both a principled estimate of the sequence of biomarker abnormality, and temporal estimates of years to symptom onset.

**Figure 7:**
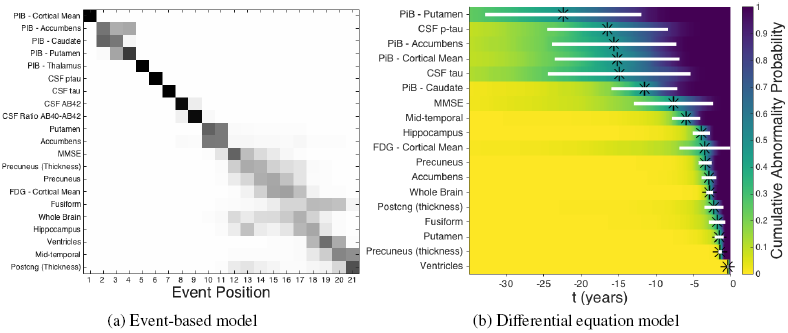
Summary: data-driven models of dominantly-inherited Alzheimer’s disease progression. (a) Event-based model for all mutation carriers in DIAN, from Figure 1. Biomarkers (imaging, molecular, cognitive) along the vertical axis are ordered by the maximum likelihood disease progression sequence (from top to bottom). The horizontal axis shows variance in the posterior sequence sampled using Markov chain Monte Carlo, with positional likelihood given by grayscale intensity. (b) Differential equation models. Each model-estimated biomarker trajectory (see Figure 4 and Supplementary Figures) estimates a probabilistic Abnormality Transition Time (years from canonical normal to canonical abnormal) and corresponding cumulative/empirical probability of abnormality (see Figure 5). Biomarkers along the vertical axis are ordered by the estimated sequence in which they reach 50% cumulative probability of abnormality (black asterisks). The viridis colour scale shows cumulative probability of abnormality increasing from the left (normal, yellow) to the right (abnormal, blue) as a function of years prior to canonical abnormality. White horizontal bars show the interquartile range of the Abnormality Transition Time density, which visualizes the rate and duration of biomarker progression.

## 4. Discussion

In this section we discuss our results further and highlight new findings that warrant further investigation. To summarise, we have reported data-driven estimates of dominantly-inherited Alzheimer’s disease progression using two modelling approaches without reliance upon familial age of onset, such as EYO, as a proxy for disease progression. The models reveal probabilistic sequences of biomarker abnormality from cross-sectional data across mutation groups, and probabilistic estimates of biomarker trajectories from a cross-section of short-term longitudinal data.

The sequences and timescales broadly agree with current understanding of dominantly-inherited Alzheimer’s disease, while producing superior detail and predictive utility than previous work.

### 4.1. Cross-sectional: event-based models

The event-based model finds a distinct ordering of biomarker abnormality events in mutation carriers: amyloid deposition measured by PiB-PET, neurofibrillary tangles and amyloid plaques in CSF, followed by a pattern of regional volume loss on MRI that is characteristic of Alzheimer’s disease, which is interspersed with declining cognitive test scores and hypometabolism measured by FDG-PET. Although the sequence shows strong agreement across different mutation types (*PSEN1*, *PSEN2*, *APP*), and APOE4 carrier groups (positive and negative), we found some small, subtle differences that warrant further investigation. For example, there was earlier abnormality in CSF A*β*42 (than CSF tau) in the APP and APOE4-positive groups, but the reverse was found in other groups. The latter could be explained by non-monotonic dynamics of CSF A*β*42 in dominantly-inherited Alzheimer’s disease (an increase followed by a decrease) as suggested by results in previous investigations (Fagan et al., 2014, Reiman et al., 2012), and consistent with our own differential equation modelling investigation (see discussion in the following subsection and Supplementary Material). Previous multimodal biomarker studies of dominantly-inherited Alzheimer’s disease (Bateman et al., 2012, Benzinger et al., 2013, Fleisher et al., 2015) are in general agreement with the event-based model sequence: amyloidosis precedes hypometabolism, neurodegeneration, and cognitive decline. Note that we considered cross-sectional volumes of brain regions, not direct measures of atrophy, which can explain why cognitive decline appears earlier than might be expected (Young et al., 2014). Importantly, all previous approaches relied upon a familial age of symptom onset as a proxy for disease progression, which intrinsically limits the accuracy of predictions due to the known imprecision in such estimates (Ryman et al., 2014). Further, such models cannot be easily generalized to sporadic forms of disease, whereas ours can.

The similarity of the event-based model sequence for dominantly-inherited Alzheimer’s disease with that for sporadic Alzheimer’s disease in previous work (Young et al., 2014) supports the notion that these two forms of Alzheimer’s disease have similar underlying disease mechanisms, and therefore that drugs developed on dominantly-inherited Alzheimer’s disease may be efficacious in sporadic Alzheimer’s disease. We note some slight deviations of the dominantly-inherited Alzheimer’s disease sequence here from the sporadic Alzheimer’s disease sequence in Young et al. (2014): the involvement of the putamen, nucleus accumbens, precuneus and posterior cingulate. Other dominantly-inherited Alzheimer’s disease investigations have observed involvement of the precuneus and cingulate regions, e.g., Benzinger et al. (2013), Cash et al. (2013), Scahill et al. (2002). Our earlier study of sporadic Alzheimer’s disease did not include these regions in the analysis, so further work will be required to determine their involvement in sporadic Alzheimer’s disease event-based models. Moreover, the nature of the biomarkers we use here means that we cannot determine whether sporadic Alzheimer’s disease and dominantly-inherited Alzheimer’s disease are similar on the microscopic scale.

The staging system provided by the event-based model has potential practical utility. In particular, it provides high classification accuracy for discriminating between pre-symptomatic and symptomatic dominantly-inherited Alzheimer’s disease mutation carriers. It correctly assigns all non-carriers to the ‘completely normal’ category (stage 0); and shows good longitudinal consistency, with event-based model stage generally increasing or remaining stable at patient follow-up. This encourages us to suggest that the staging system has utility in future clinical trials, both for screening of potential participants and for defining end-points. For example, recruiting individuals at event-based model stages 1–5 (Figure 1: PiB-PET abnormality only), and defining an end-point as reaching stage 8 (addition of CSF abnormality). The same approach could work for personalised treatment assignment. For example, an anti-amyloid agent might only be appropriate for APOE4-positive individuals at event-based model stages 1 and 2 (Figure 2(a)).

We found the event-based model stages to correlate strongly with cognitive status: cognitively normal participants were assigned early model stages, symptomatic dominantly-inherited Alzheimer’s disease participants were assigned late model stages, and participants with mild symptoms were more spread out across the stages. The mildly symptomatic group in dominantly-inherited Alzheimer’s disease were the most heterogeneous, which is in agreement with our results in sporadic Alzheimer’s disease (Young et al., 2014), but possibly for different reasons. One contributing factor in dominantly-inherited Alzheimer’s disease is that the mildly symptomatic group may include unaffected mutation carriers whose anxiety about their mutation status manifested as apparent cognitive abnormality and contributed to their diagnosis (global CDR = 0·5). In any case, the fine-grained disease staging offered by the event-based model can shed light upon the heterogeneity contained within a prodromal disease stage. Separate work will consider explicitly modelling prodromal disease phases within the event-based model. Supplementary Figure 15(a) shows that event-based model stage correlates with familial age of onset, although further follow-up will be required to ascertain the predictive utility of event-based model stage compared to familial age of onset by looking at a large number of individuals who develop clinical Alzheimer’s disease dementia during a study of dominantly-inherited Alzheimer’s disease.

### 4.2. Longitudinal: differential equation models

Our nonparametric fits to differential biomarker data are data-driven probabilistic estimates of an underlying differential equation driving the disease biomarker evolution. This approach is capable of estimating monotonic biomarker trajectories only because the absence of a known disease time precludes estimation of non-monotonic trajectories such as those found in the cross-sectional analysis of CSF tau and p-tau reported in Fagan et al. (2014). Further, the use of a single differential data point per participant precludes modelling within-individual dynamics using this approach. The consequence is that if enough individuals display contrary dynamics to the average, perhaps due to measurement noise for example, then a sensible trajectory cannot be inferred. This happened for CSF markers of tau, A*β*42, and the A*β*40/A*β*42 ratio, as shown in Supplementary Figure 14. Otherwise, we obtained trajectory estimates for the same set of biomarkers in the event-based model results (Figure 4 and Supplementary Material). Most differential-equation-model-estimated biomarker trajectories showed accelerating dynamics, with little or no apparent deceleration, which may arise from under-sampling of later disease stages (for example because DIAN recruitment was focussed on pre-symptomatic dominantly-inherited Alzheimer’s disease). The magenta fits in Figure 4 correspond to those in Bateman et al. (2012) (taken directly from the supplementary material in that paper), which was a cross-sectional regression of biomarker trends as a function of EYO, in the DIAN dataset. It is apparent from Figure 4 that the most noteworthy difference between the differential equation model trajectories and EYO regression trajectories are the slower post-onset dynamics estimated for hippocampal volume when using the latter. The cross-sectional approach, such as in Bateman et al. (2012) and Benzinger et al. (2013), is less able to capture speed of progression than the differential equation modelling approach, which utilises short-duration longitudinal data, within subjects. This is supported by the longitudinal analysis in the EYO-based regional imaging biomarker investigation in Benzinger et al. (2013), which found that the cross-sectional biomarker trajectory tended to underestimate the slope of individual trajectories, post-onset.

We did not model biomarker measurement noise. Such noise can lead to regression dilution, which, in a differential equation modelling approach, would produce an elongated (slower) biomarker trajectory. Thus, temporal quantities we have estimated, such as abnormality transition times, may represent overestimates — particularly for biomarkers with large measurement noise. However, regression dilution was probably not a problem, as evidenced by our ability to accurately predict actual symptom onset, which we discuss further in the following paragraph.

Recently, Ryman et al. (2014) performed a meta-analysis of actual symptom onset in multiple studies of dominantly-inherited Alzheimer’s disease including DIAN, and considered prediction of age at symptom onset using ages of onset for parents, family average, and group-wise averages by mutation type, as well as APOE4 genotype and sex. They argued that mutation type and family history should be used to estimate onset in clinical research. This conclusion was reached by analysing the proportion of variance in actual age of onset that could be explained by these factors in a linear regression scenario, quantified by adjusted *r*^2^. Specifically, they found *r*^2^ = 0·3838 (parental), *r*^2^ = 0·4906 (family average) and *r*^2^ = 0·5225 (mutation type). For clinical utility we argue that a model’s predictive accuracy should be quantified, such as by using root-mean-squared error in prediction of unseen data. We quantified predictive accuracy for six participants in the DIAN dataset with observed symptom onset (at Data Freeze 6) in Figure 6 — we found root-mean-squared error of 5·54 years with *r*^2^ = 0.37 (parental), and root-mean-squared error of 8·61 years with *r*^2^ = 0·33 (mutation type), whereas our data-driven model-based approach performed considerably better: root-mean-squared error of 1·35 years with *r*^2^ = 0·44.

### 4.3. Summary

Dominantly-inherited Alzheimer’s disease progression occurs over multiple decades. Our two data-driven approaches have estimated dominantly-inherited Alzheimer’s disease progression models by combining shorter cross-sections of data. This was made possible in part by assuming a single progression pattern across individuals. Despite this, our models are able to predict probabilistic outcomes for individuals by comparing them to the average pattern. With increased availability of data, especially actual symptom onset, an important future aim is to incorporate multilevel modelling to improve the specificity of predictions across mutation types, families, and individuals, and to hopefully understand more of the heterogeneity observed in dominantly-inherited Alzheimer’s disease.

Our probabilistic, EYO-agnostic, data-driven computational models of dominantly-inherited Alzheimer’s disease reveal evidence-based patterns in the progression of this relatively rare disease. The similarities with sporadic Alzheimer’s disease progression provides encouragement for ongoing trials into anti-amyloid therapies such as the ones currently underway by the DIAN Trials Unit. We have also demonstrated abilities of the data-driven models for fine-grained patient staging and prognosis, which promises utility for recruitment, stratification, and surrogate outcome measures in clinical trials.

## Acknowledgements

Data collection and sharing for this project was supported by The Dominantly Inherited Alzheimer Network (DIAN; UF1 AG032438; to RJB and JCM), funded by the National Institute on Aging, the German Center for Neurodegenerative Diseases, the Medical Research Council (MRC; to NCF) Dementias Platform UK (MR/L023784/1 and MR/009076/1), and National Institute for Health Research Queen Square Dementia Biomedical Research Unit. JCM and RJB receive research support from National Institute of Health. RJB receives research support from the Alzheimer’s Association, Foundation for Biomedical Research and Innovation, BrightFocus Foundation, Cure Alzheimer’s Fund, Glenn Foundation for Medical Research, Metropolitan Life Foundation, and Ruth K Broadman Biomedical Research Foundation. This manuscript has been reviewed by DIAN Study investigators for scientific content and consistency of data interpretation with previous DIAN Study publications. We gratefully acknowledge the altruism of the participants and their families and contributions of the DIAN research and support staff at each of the participating sites for their contributions to this study. The DIAN Expanded Registry welcomes contact from any families or treating clinicians interested in research about dominantly inherited familial Alzheimer’s disease.

## Supplementary Material

Supplementary Figure 1 shows cross-validation of the event-based models in Figure 2. Differential equation model fits for selected biomarkers in the DIAN dataset which displayed monotonic behaviour on average are spread between three figures, for aesthetic reasons:

- Supplementary Figure 2 (cross-validation in Supplementary Figure 8);
- Supplementary Figure 3 (cross-validation in Supplementary Figure 9);
- Supplementary Figure 4 (cross-validation in Supplementary Figure 10).

Corresponding biomarker trajectories are shown in:

- Supplementary Figure 5 (cross-validation in Supplementary Figure 11);
- Supplementary Figure 6 (cross-validation in Supplementary Figure 12);
- Supplementary Figure 7 (cross-validation in Supplementary Figure 13).

Supplementary Table 1 is a numerical summary of the differential equation model fitting results: the model hyperparameter estimates and numerical convergence of the Markov chain Monte Carlo fits via the potential scale reduction factor (Gelman et al., 2014, Vehtari et al., 2016). Ten-fold cross-validation results are quoted as mean ± standard deviation across the ten folds.

Supplementary Figure 14 shows differential equation model fits for the few CSF amyloid biomarkers where the approach was incapable of estimating valid biomarker trajectories due apparently to non-monotonic dynamics.

Supplementary Figure 15 compares our model-based measures of dominantly-inherited Alzheimer’s disease progression to the traditional measure, EYO from parental age of onset. Further discussion of this is included below.

**Supplementary Table 1:**
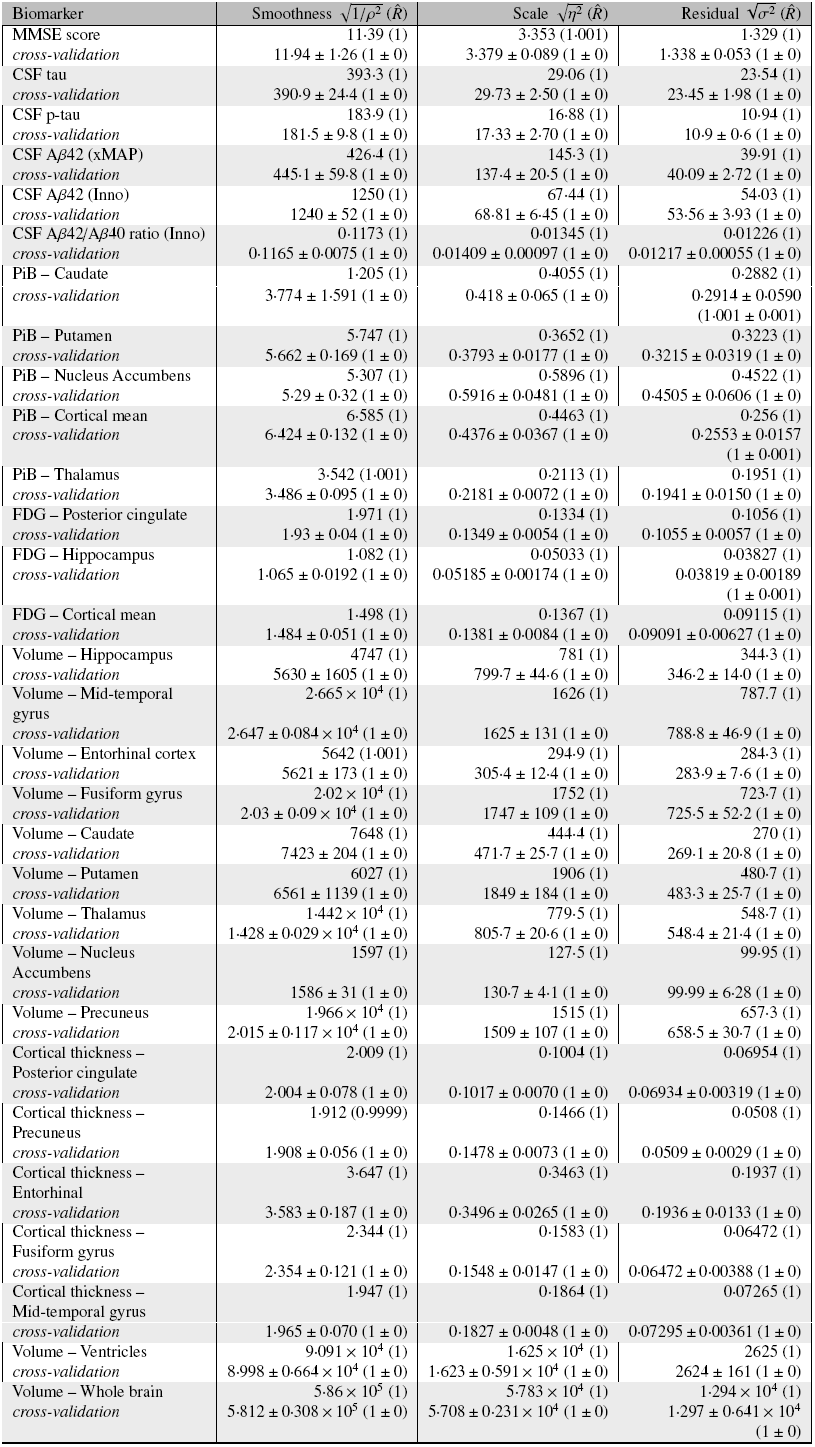
Differential equation model regression results: Gaussian process hyperparameter estimates. Units — CSF: pg/mL (except ratio); PET (PiB and FDG): standardized uptake value ratio (SUVR) relative to the cerebellum; Volumes: mm^3^; Cortical Thickness (CT): mm. Ten-fold cross validation shows mean ± standard deviation. The potential scale reduction factor 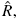, here given precise to three decimal places, indicates convergence of the algorithm: values close to 1 indicate strong convergence.

| Biomarker | Smoothness $\sqrt{1/\rho^2} (\hat{R})$ | Scale $\sqrt{\eta^2} (\hat{R})$ | Residual $\sqrt{\sigma^2} (\hat{R})$ |
| --- | --- | --- | --- |
| MMSE score | 11.39 (1) | 3.353 (1.001) | 1.329 (1) |
| *cross-validation* | 11.94 ± 1.26 (1 ± 0) | 3.379 ± 0.089 (1 ± 0) | 1.338 ± 0.053 (1 ± 0) |
| CSF tau | 393.3 (1) | 29.06 (1) | 23.54 (1) |
| *cross-validation* | 390.9 ± 24.4 (1 ± 0) | 29.73 ± 2.50 (1 ± 0) | 23.45 ± 1.98 (1 ± 0) |
| CSF p-tau | 183.9 (1) | 16.88 (1) | 10.94 (1) |
| *cross-validation* | 181.5 ± 9.8 (1 ± 0) | 17.33 ± 2.70 (1 ± 0) | 10.9 ± 0.6 (1 ± 0) |
| CSF Aβ42 (xMAP) | 426.4 (1) | 145.3 (1) | 39.91 (1) |
| *cross-validation* | 445.1 ± 59.8 (1 ± 0) | 137.4 ± 20.5 (1 ± 0) | 40.09 ± 2.72 (1 ± 0) |
| CSF Aβ42 (Inno) | 1250 (1) | 67.44 (1) | 54.03 (1) |
| *cross-validation* | 1240 ± 52 (1 ± 0) | 68.81 ± 6.45 (1 ± 0) | 53.56 ± 3.93 (1 ± 0) |
| CSF Aβ42/Aβ40 ratio (Inno) | 0.1173 (1) | 0.01345 (1) | 0.01226 (1) |
| *cross-validation* | 0.1165 ± 0.0075 (1 ± 0) | 0.01409 ± 0.00097 (1 ± 0) | 0.01217 ± 0.00055 (1 ± 0) |
| PiB – Caudate | 1.205 (1) | 0.4055 (1) | 0.2882 (1) |
| *cross-validation* | 3.774 ± 1.591 (1 ± 0) | 0.418 ± 0.065 (1 ± 0) | 0.2914 ± 0.0590<br>(1.001 ± 0.001) |
| PiB – Putamen | 5.747 (1) | 0.3652 (1) | 0.3223 (1) |
| *cross-validation* | 5.662 ± 0.169 (1 ± 0) | 0.3793 ± 0.0177 (1 ± 0) | 0.3215 ± 0.0319 (1 ± 0) |
| PiB – Nucleus Accumbens | 5.307 (1) | 0.5896 (1) | 0.4522 (1) |
| *cross-validation* | 5.29 ± 0.32 (1 ± 0) | 0.5916 ± 0.0481 (1 ± 0) | 0.4505 ± 0.0606 (1 ± 0) |
| PiB – Cortical mean | 6.585 (1) | 0.4463 (1) | 0.256 (1) |
| *cross-validation* | 6.424 ± 0.132 (1 ± 0) | 0.4376 ± 0.0367 (1 ± 0) | 0.2553 ± 0.0157<br>(1 ± 0.001) |
| PiB – Thalamus | 3.542 (1.001) | 0.2113 (1) | 0.1951 (1) |
| *cross-validation* | 3.486 ± 0.095 (1 ± 0) | 0.2181 ± 0.0072 (1 ± 0) | 0.1941 ± 0.0150 (1 ± 0) |
| FDG – Posterior cingulate | 1.971 (1) | 0.1334 (1) | 0.1056 (1) |
| *cross-validation* | 1.93 ± 0.04 (1 ± 0) | 0.1349 ± 0.0054 (1 ± 0) | 0.1055 ± 0.0057 (1 ± 0) |
| FDG – Hippocampus | 1.082 (1) | 0.05033 (1) | 0.03827 (1) |
| *cross-validation* | 1.065 ± 0.0192 (1 ± 0) | 0.05185 ± 0.00174 (1 ± 0) | 0.03819 ± 0.00189<br>(1 ± 0.001) |
| FDG – Cortical mean | 1.498 (1) | 0.1367 (1) | 0.09115 (1) |
| *cross-validation* | 1.484 ± 0.051 (1 ± 0) | 0.1381 ± 0.0084 (1 ± 0) | 0.09091 ± 0.00627 (1 ± 0) |
| Volume – Hippocampus | 4747 (1) | 781 (1) | 344.3 (1) |
| *cross-validation* | 5630 ± 1605 (1 ± 0) | 799.7 ± 44.6 (1 ± 0) | 346.2 ± 14.0 (1 ± 0) |
| Volume – Mid-temporal gyrus | 2.665 × 10<sup>4</sup> (1) | 1626 (1) | 787.7 (1) |
| *cross-validation* | 2.647 ± 0.084 × 10<sup>4</sup> (1 ± 0) | 1625 ± 131 (1 ± 0) | 788.8 ± 46.9 (1 ± 0) |
| Volume – Entorhinal cortex | 5642 (1.001) | 294.9 (1) | 284.3 (1) |
| *cross-validation* | 5621 ± 173 (1 ± 0) | 305.4 ± 12.4 (1 ± 0) | 283.9 ± 7.6 (1 ± 0) |
| Volume – Fusiform gyrus | 2.02 × 10<sup>4</sup> (1) | 1752 (1) | 723.7 (1) |
| *cross-validation* | 2.03 ± 0.09 × 10<sup>4</sup> (1 ± 0) | 1747 ± 109 (1 ± 0) | 725.5 ± 52.2 (1 ± 0) |
| Volume – Caudate | 7648 (1) | 444.4 (1) | 270 (1) |
| *cross-validation* | 7423 ± 204 (1 ± 0) | 471.7 ± 25.7 (1 ± 0) | 269.1 ± 20.8 (1 ± 0) |
| Volume – Putamen | 6027 (1) | 1906 (1) | 480.7 (1) |
| *cross-validation* | 6561 ± 1139 (1 ± 0) | 1849 ± 184 (1 ± 0) | 483.3 ± 25.7 (1 ± 0) |
| Volume – Thalamus | 1.442 × 10<sup>4</sup> (1) | 779.5 (1) | 548.7 (1) |
| *cross-validation* | 1.428 ± 0.029 × 10<sup>4</sup> (1 ± 0) | 805.7 ± 20.6 (1 ± 0) | 548.4 ± 21.4 (1 ± 0) |
| Volume – Nucleus Accumbens | 1597 (1) | 127.5 (1) | 99.95 (1) |
| *cross-validation* | 1586 ± 31 (1 ± 0) | 130.7 ± 4.1 (1 ± 0) | 99.99 ± 6.28 (1 ± 0) |
| Volume – Precuneus | 1.966 × 10<sup>4</sup> (1) | 1515 (1) | 657.3 (1) |
| *cross-validation* | 2.015 ± 0.117 × 10<sup>4</sup> (1 ± 0) | 1509 ± 107 (1 ± 0) | 658.5 ± 30.7 (1 ± 0) |
| Cortical thickness – Posterior cingulate | 2.009 (1) | 0.1004 (1) | 0.06954 (1) |
| *cross-validation* | 2.004 ± 0.078 (1 ± 0) | 0.1017 ± 0.0070 (1 ± 0) | 0.06934 ± 0.00319 (1 ± 0) |
| Cortical thickness – Precuneus | 1.912 (0.9999) | 0.1466 (1) | 0.0508 (1) |
| *cross-validation* | 1.908 ± 0.056 (1 ± 0) | 0.1478 ± 0.0073 (1 ± 0) | 0.0509 ± 0.0029 (1 ± 0) |
| Cortical thickness – Entorhinal | 3.647 (1) | 0.3463 (1) | 0.1937 (1) |
| *cross-validation* | 3.583 ± 0.187 (1 ± 0) | 0.3496 ± 0.0265 (1 ± 0) | 0.1936 ± 0.0133 (1 ± 0) |
| Cortical thickness – Fusiform gyrus | 2.344 (1) | 0.1583 (1) | 0.06472 (1) |
| *cross-validation* | 2.354 ± 0.121 (1 ± 0) | 0.1548 ± 0.0147 (1 ± 0) | 0.06472 ± 0.00388 (1 ± 0) |
| Cortical thickness – Mid-temporal gyrus | 1.947 (1) | 0.1864 (1) | 0.07265 (1) |
| *cross-validation* | $1.965 \pm 0.070 (1 \pm 0)$ | $0.1827 \pm 0.0048 (1 \pm 0)$ | $0.07295 \pm 0.00361 (1 \pm 0)$ |
| Volume – Ventricles | $9.091 \times 10^4 (1)$ | $1.625 \times 10^4 (1)$ | 2625 (1) |
| *cross-validation* | $8.998 \pm 0.664 \times 10^4 (1 \pm 0)$ | $1.623 \pm 0.591 \times 10^4 (1 \pm 0)$ | $2624 \pm 161 (1 \pm 0)$ |
| Volume – Whole brain | $5.86 \times 10^5 (1)$ | $5.783 \times 10^4 (1)$ | $1.294 \times 10^4 (1)$ |
| *cross-validation* | $5.812 \pm 0.308 \times 10^5 (1 \pm 0)$ | $5.708 \pm 0.231 \times 10^4 (1 \pm 0)$ | $1.297 \pm 0.641 \times 10^4 (1 \pm 0)$ |

Figure 15 shows correspondence between different estimates of disease progression: (a) event-based model and EYO; (b) ETO and EYO; and (c) event-based model and ETO. In these figures, disease progresses from left to right and from bottom to top. Broadly speaking, Figure 15 shows that all estimates of disease progression correctly predict unaffected/asymptomatic individuals (global CDR = 0) to lie towards the lower left corner and affected individuals (CDR > 0) towards the upper right. Further, there is quite good linear correlation across all estimates, as shown by the linear regression fits in the left column, and by the quantile plots in the right column. In Figure 15(a), unaffected individuals should be found towards the lower left corner (EYO < 0; early event-based model stage), with affected individuals towards the top right corner (EYO > 0; late event-based model stage). Indeed, this is what we see, with good overall correlation between EYO and event-based model stage across diagnostic groups. For all groups of mutation carriers combined, a linear fit yields *r*^2^ ≈ 0·45. Figure 15(b) shows a linear relationship between ETO and EYO for all mutation carriers in the DIAN dataset: both symptomatic and asymptomatic individuals. Figure 15(c) compares our two modelling approaches by plotting ETO against event-based model stage.

**Supplementary Figure 1:**
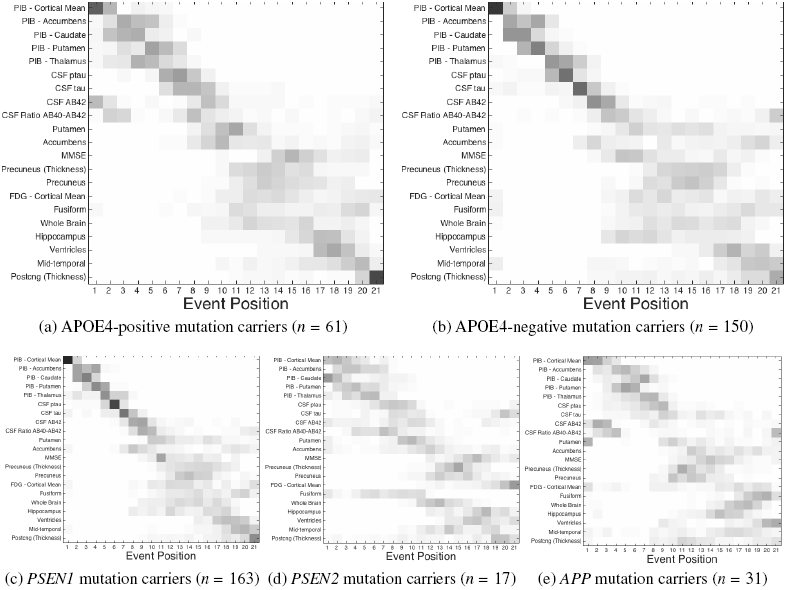
Cross-validation of event-based models of dominantly-inherited Alzheimer’s disease: separated into mutation groups (see Figure 2).

**Supplementary Figure 2:**
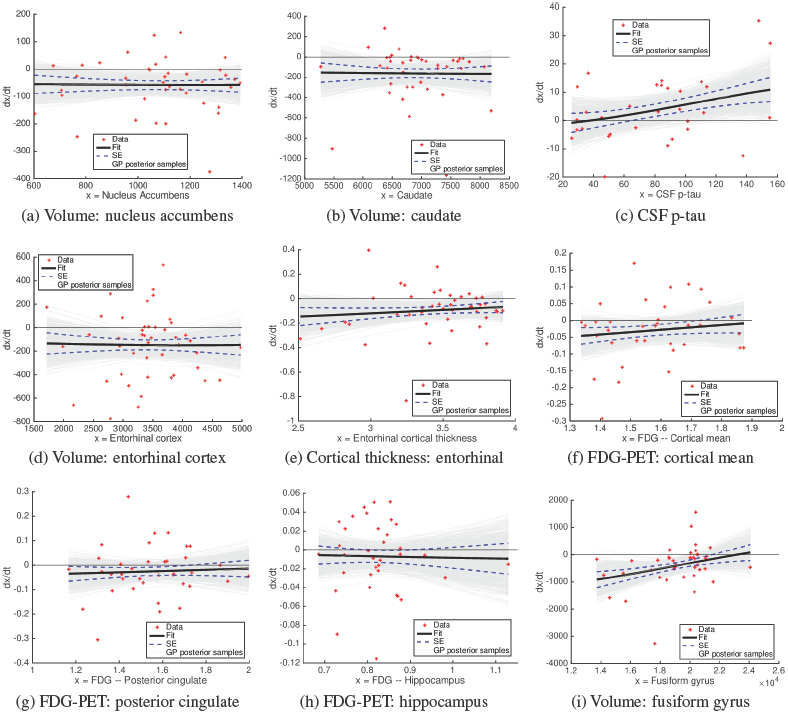
Differential equation model fits (1 of 3). Data points are shown as red plusses. The Bayesian nonparametric differential regression fits are shown as mean (heavy solid black line) ± standard error (SE, dashed blue lines), and samples from the full posterior (light grey lines). Gaussian process model hyperparameters are in Supplementary Table 1. Corresponding biomarker trajectories are shown in Supplementary Figure 5.

**Supplementary Figure 3:**
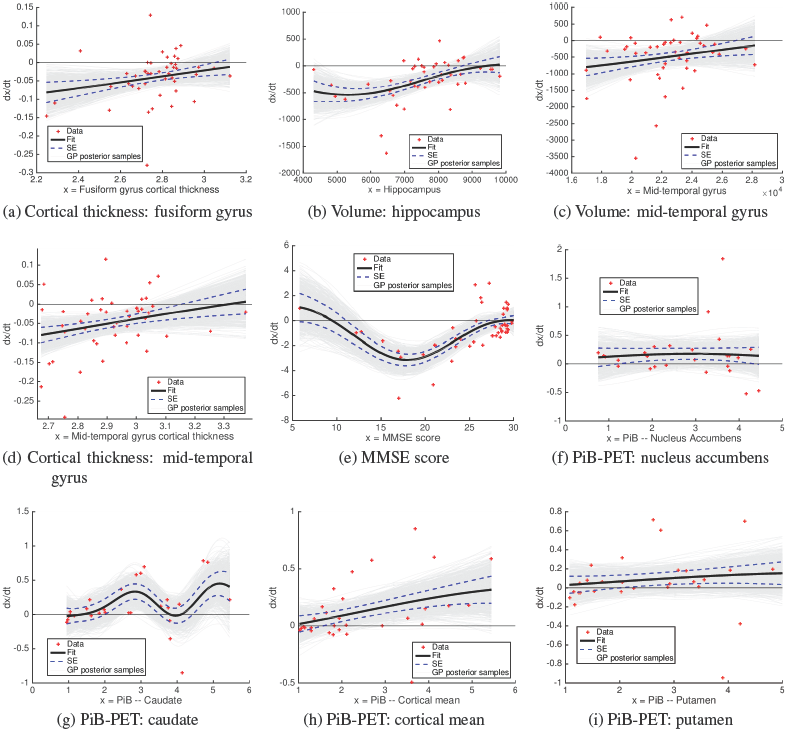
Differential equation model fits (2 of 3). Corresponding biomarker trajectories shown in Supplementary Figure 6. Key explained in Supplementary Figure 2.

**Supplementary Figure 4:**
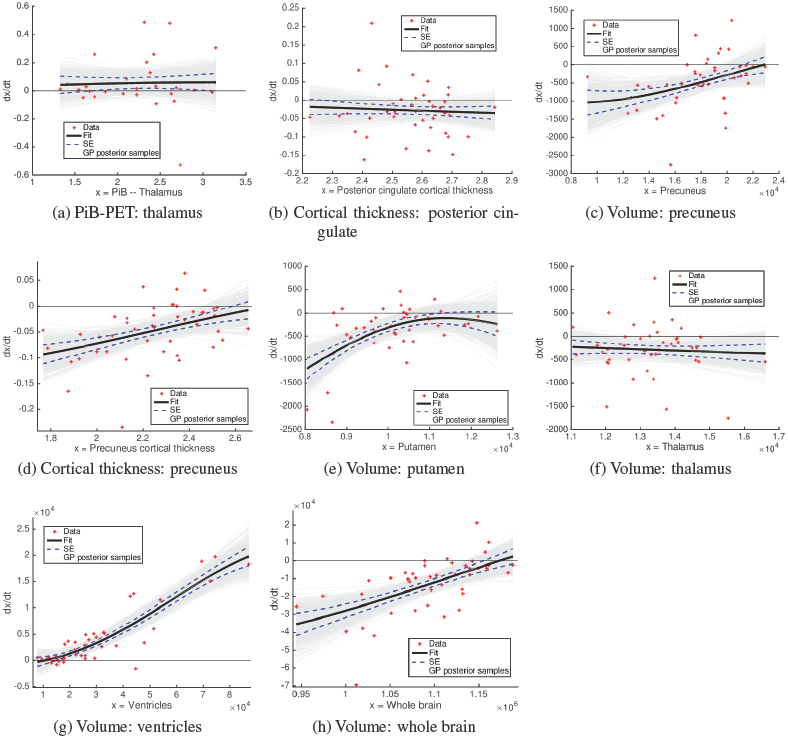
Differential equation model fits (3 of 3). Corresponding biomarker trajectories shown in Supplementary Figure 7. Key explained in Supplementary Figure 2.

**Supplementary Figure 5:**
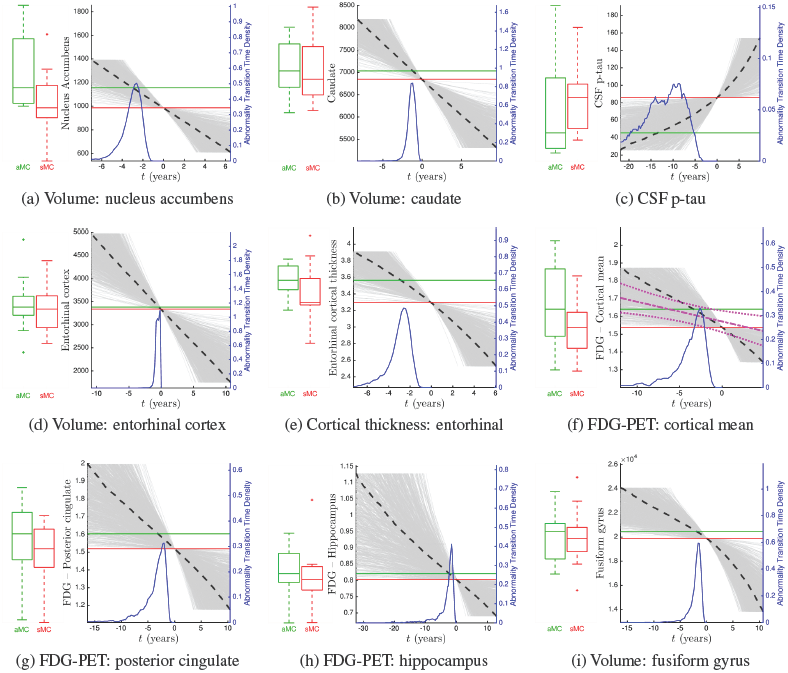
Biomarker trajectories (1 of 3). Corresponding differential equation model fits shown in Supplementary Figure 2. Key explained in Figure 4.

**Supplementary Figure 6:**
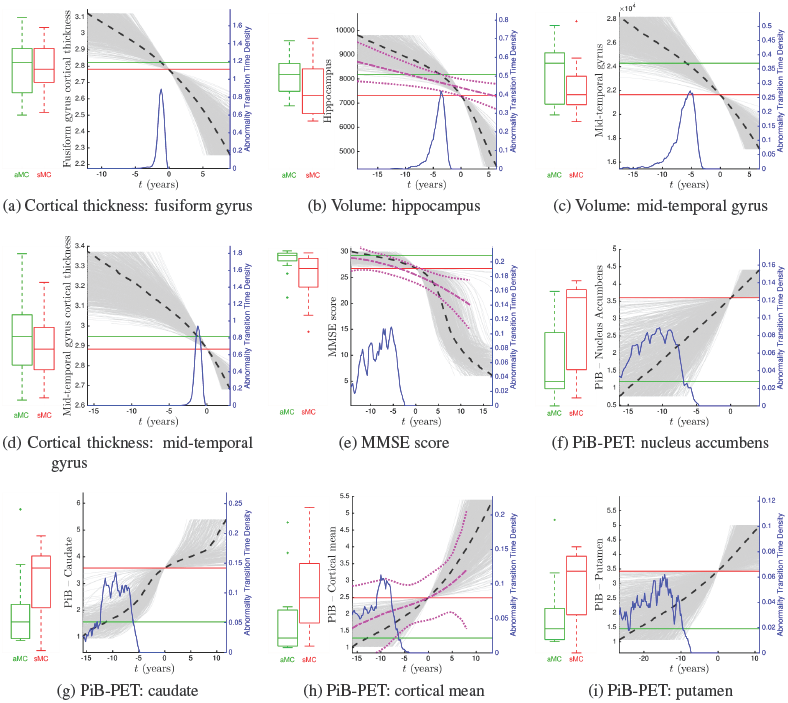
Biomarker trajectories (2 of 3). Corresponding differential equation model fits shown in Supplementary Figure 3. Key explained in Figure 4.

**Supplementary Figure 7:**
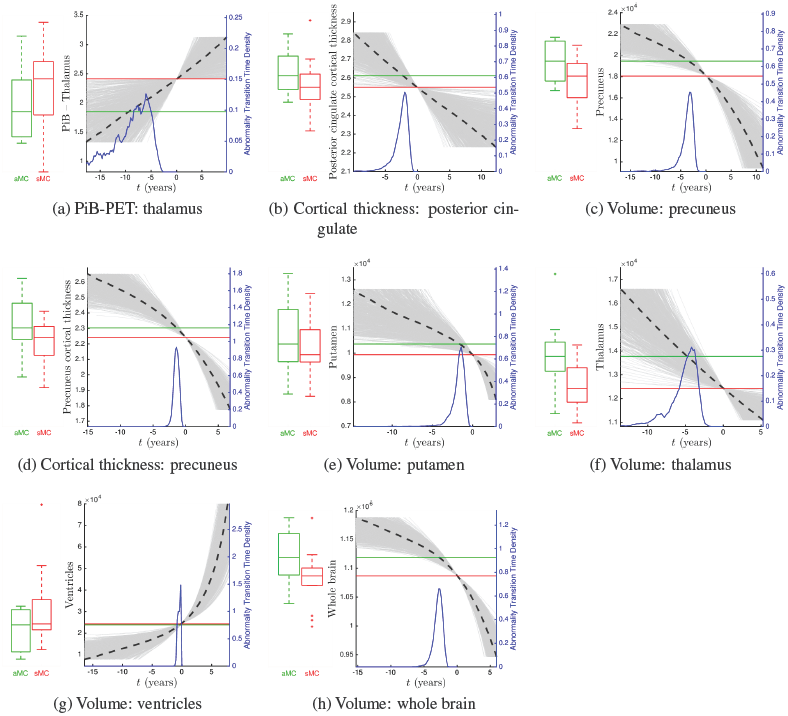
Biomarker trajectories (3 of 3). Corresponding differential equation model fits shown in Supplementary Figure 4. Key explained in Figure 4.

**Supplementary Figure 8:**
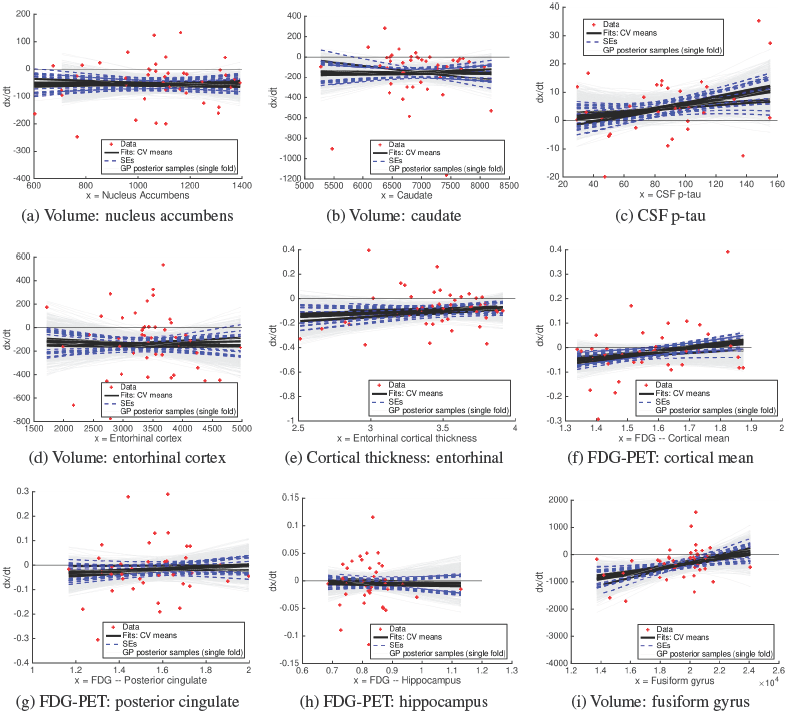
Differential equation model fits: ten-fold cross-validation (1 of 3) of the fits in Supplementary Figure 2. Corresponding biomarker trajectories are in Supplementary Figure 11.

**Supplementary Figure 9:**
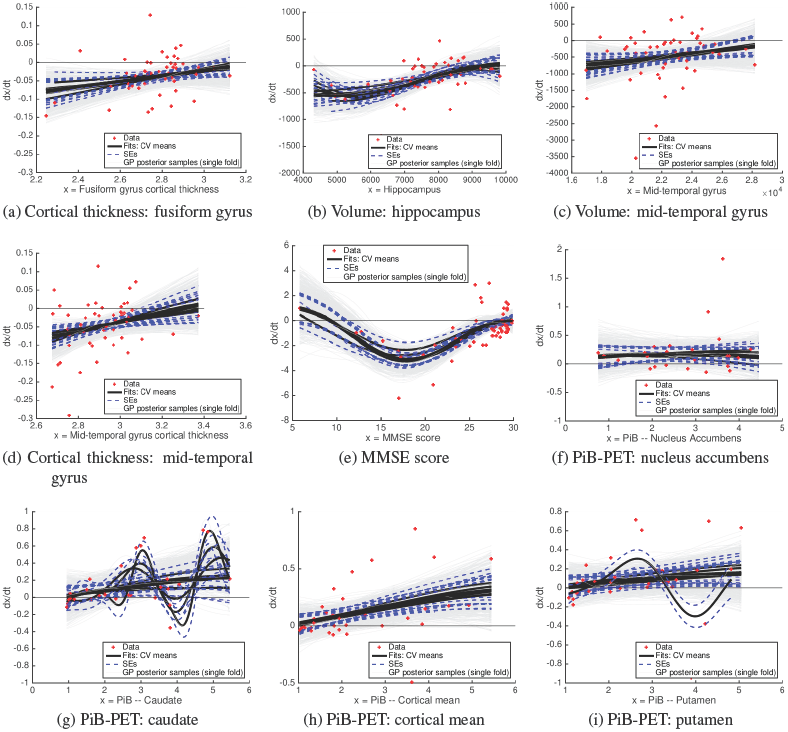
Differential equation model fits: ten-fold cross-validation (2 of 3) of the fits in Supplementary Figure 3. Corresponding biomarker trajectories are in Supplementary Figure 12.

**Supplementary Figure 10:**
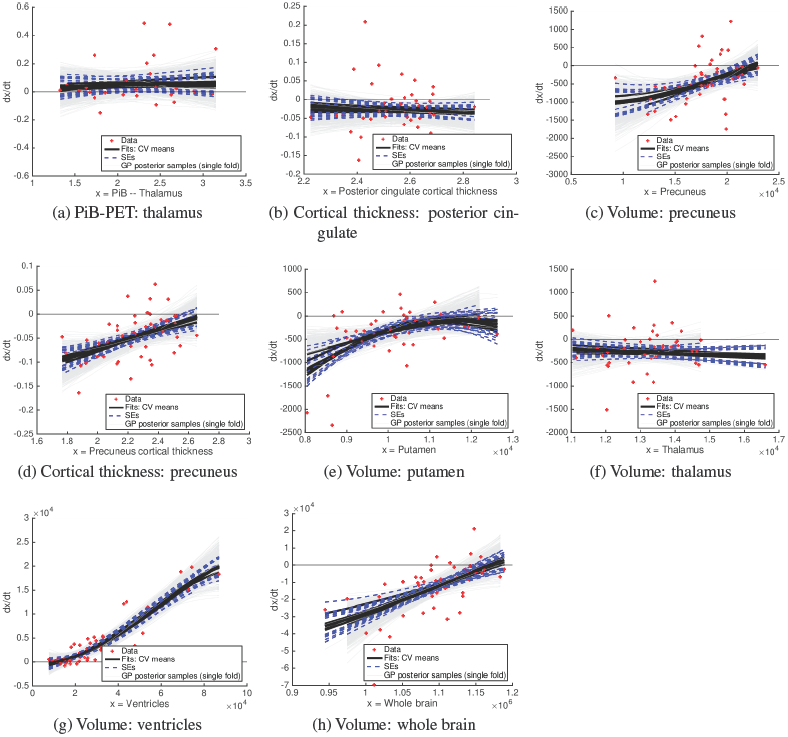
Differential equation model fits: ten-fold cross-validation (3 of 3) of the fits in Supplementary Figure 4. Corresponding biomarker trajectories are in Supplementary Figure 13.

**Supplementary Figure 11:**
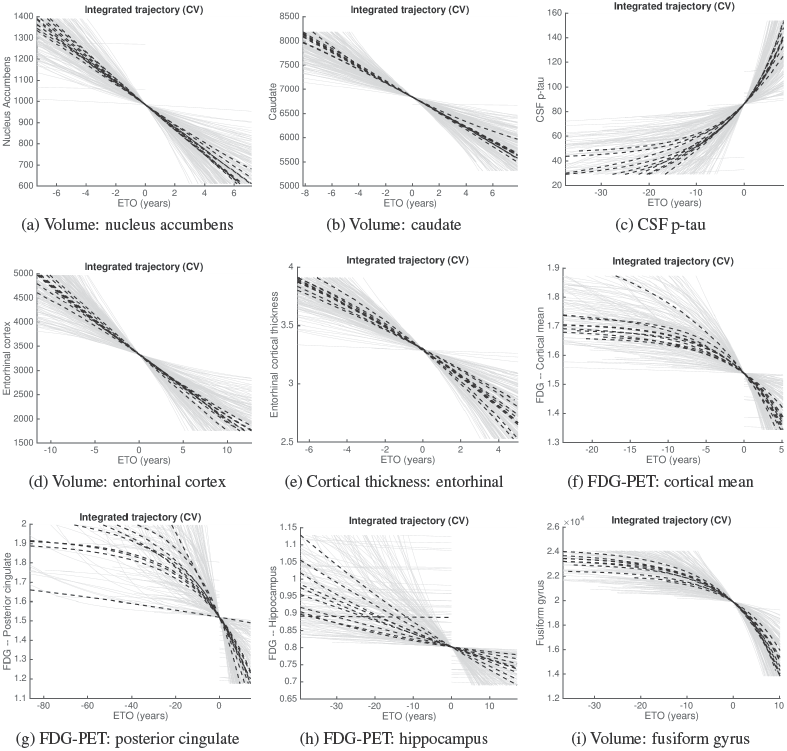
Biomarker trajectories: ten-fold cross-validation (1 of 3). Corresponding differential equation model fits are in Supplementary Figure 8.

**Supplementary Figure 12:**
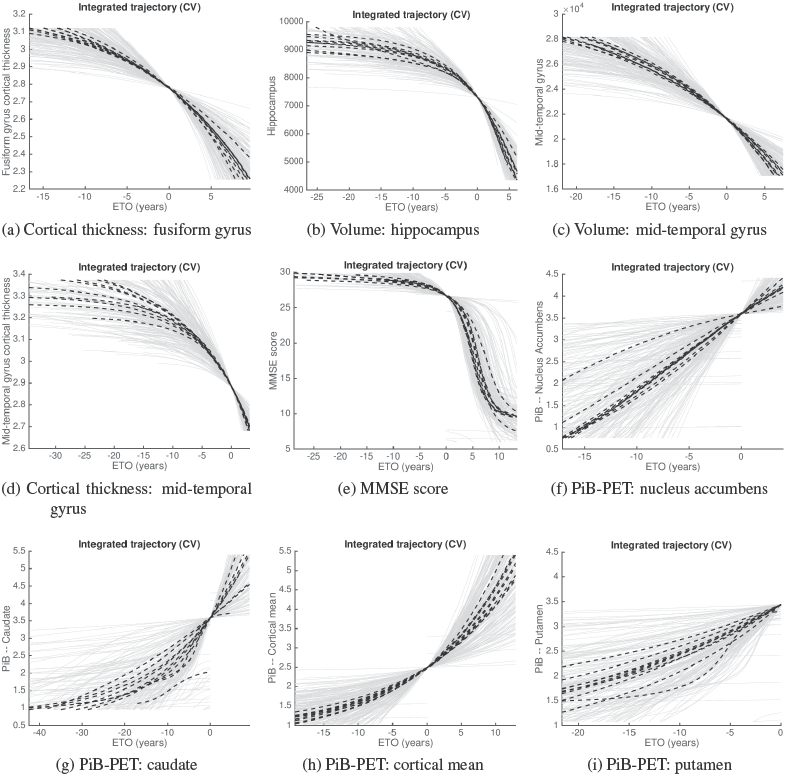
Biomarker trajectories: ten-fold cross-validation (2 of 3). Corresponding differential equation model fits are in Supplementary Figure 9.

**Supplementary Figure 13:**
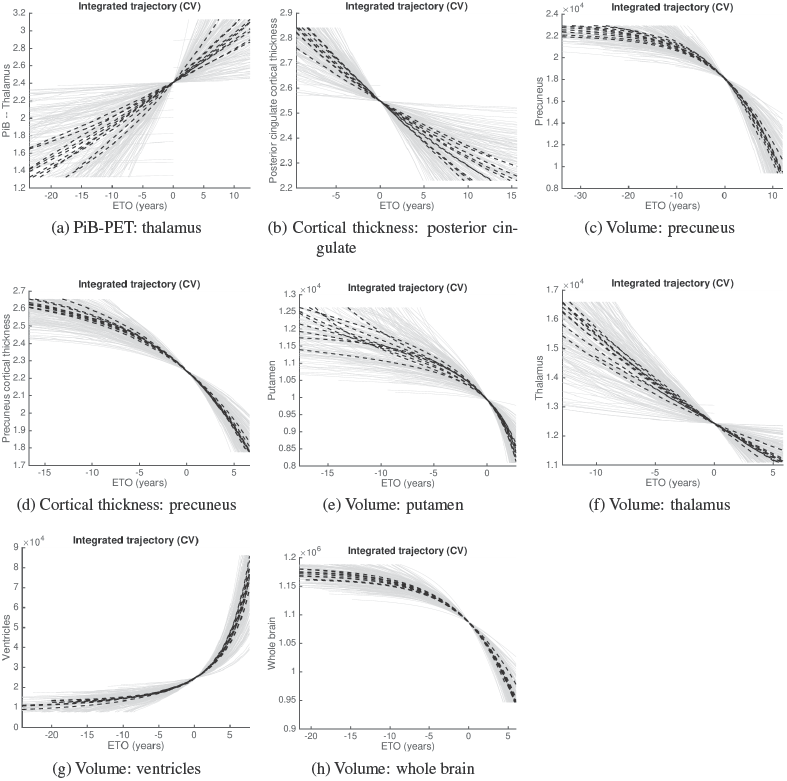
Biomarker trajectories: ten-fold cross-validation (3 of 3). Corresponding differential equation model fits are in Supplementary Figure 10.

**Supplementary Figure 14:**
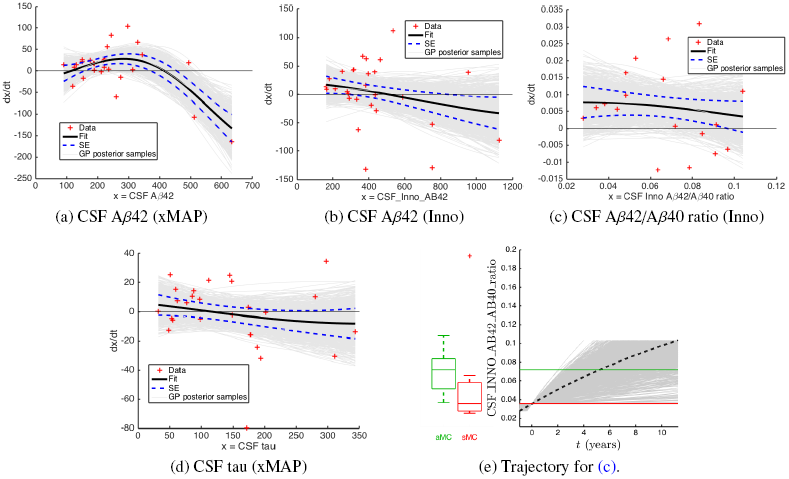
Differential equation model fits for biomarkers where the approach could not infer a valid biomarker trajectory. In (a), (b), and (d), the non-monotonic dynamics precludes inference of a single trajectory. In (c), the fit implies an increasing biomarker (on average), but this biomarker is observed in (e) to decrease as disease progresses (Trajectory for C.).

**Supplementary Figure 15:**
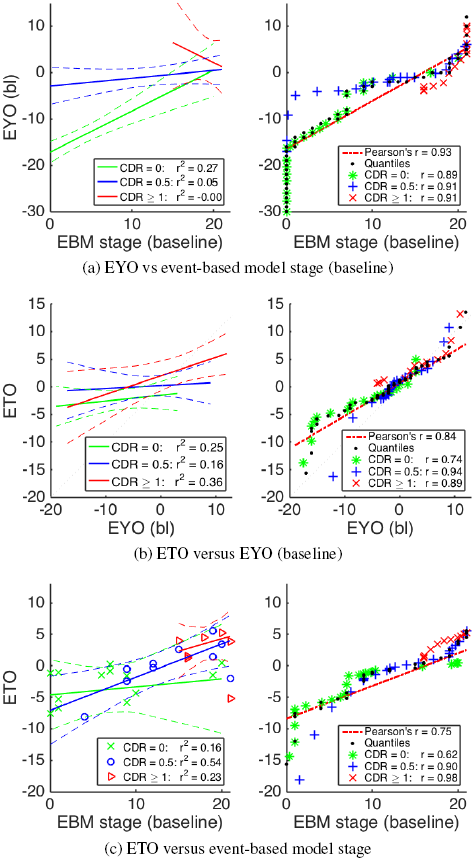
Comparison of disease progression estimates for mutation carriers in the DIAN dataset. Clinical progression is defined using global Clinical Dementia Rating (CDR) score: asymptomatic (CDR = 0), mild symptomatic (CDR = 0·5) and symptomatic (CDR > 0·5). Left panels: direct comparison of linear regression fits and 95% confidence intervals, with individual data points for EYO removed to avoid unblinding. Right panels: comparison of distributions via quantile plots, with a reference line (correlation) shown for all diagnostic categories combined. EYO = Estimated Years to Onset from parental age of onset; EBM = event-based model; ETO = Estimated Time to Onset from differential equation models; CDR = Clinical Dementia Rating; bl = baseline.

